# High-order enhancer hubs buffer allelic regulatory variation through kinetic compensation

**DOI:** 10.64898/2026.09.14.750771

**Authors:** Jiang Tan, Monica F Sentmanat, Yuqing Wu, Cody Peng, Catrina Fronick, Christopher Markovic, Xiaoxia Cui, Robert S Fulton, Richard Head, Ting Wang, Yidan Sun

## Abstract

Diploid genomes carry millions of heterozygous variants in cis-regulatory DNA, yet most genes produce similar RNA output from two parental alleles. How this balance is maintained is unclear. We developed Nanopore-HiChIP, a long-read method that maps high-order enhancer hubs on each haplotype. Over half of these enhancer hubs differ in chromatin architecture and transcription-factor occupancy between homologous chromosomes, but their target genes show substantially lower rates of allele-specific expression than genes lacking hub regulation. Single-cell kinetic modeling shows that burst frequency and burst size change in opposite directions, thereby preserving balanced transcriptional output. This hub-mediated kinetic buffering is enriched at haploinsufficient genes and coincides with smaller effects of expression quantitative trait loci. Enhancer hubs therefore absorb allelic regulatory variation through kinetic compensation, protecting dosage-sensitive transcription.

## Main text

Precise control of balanced transcription between the two parental alleles is a fundamental constraint of diploid gene regulation. For many genes, particularly haploinsufficient genes, reduced expression from a single allele is sufficient to disrupt normal development and cause human disease, highlighting the importance of preserving proper allelic dosage (*1*–*6*). Yet every diploid genome harbors millions of naturally occurring sequence differences between maternal and paternal chromosomes, many of which fall within cis-regulatory elements and can alter transcription factor binding, chromatin accessibility, and enhancer activity (*6*–*11*). These variations can therefore create substantially different regulatory states between the two parental chromosomes. Nevertheless, despite this pervasive cis-regulatory divergence, most genes maintain remarkably similar expression from their two alleles, and allele-specific expression is observed for only a small fraction of genes across mammalian genomes in a given context (*12*–*17*). How two genetically distinct parental chromosomes maintain balanced transcription despite extensive differences in their regulatory landscapes remains one of the fundamental unanswered questions in gene regulation.

One hypothesis is that gene regulatory architecture buffers local perturbation. Enhancers frequently act in groups rather than in isolation, and multiple enhancers can regulate the same gene with partially redundant activities (*18*–*21*). Genetic perturbation studies have shown that disruption of individual enhancers often produces surprisingly modest effects on gene expression, suggesting that enhancer redundancy contributes to transcriptional robustness (*22*–*25*). Similarly, population genetic studies have reported that genes associated with redundant enhancer domains are depleted of detectable cis-eQTLs despite being enriched for disease-associated genes (*26*), implying that enhancer redundancy may buffer the transcriptional consequences of regulatory variation. However, these observations remain largely correlative, and the molecular mechanism by which enhancer redundancy preserves balanced transcription has remained unknown.

This gap has persisted due to technical limitations. Current enhancer-focused chromosome conformation technologies, such as promoter capture Hi-C (*27*), HiChIP/PLAC-seq (*28*, *29*), and HiCAR (*30*), primarily resolve pairwise chromatin interactions and cannot reconstruct the high-order organization of multiple enhancers acting together on the same promoter. Although recent approaches like GAM, SPRITE, and Pore-C (*31*–*35*) recover multi-way chromatin contacts, they are not selective for active regulatory interactions, making it difficult to resolve enhancer hubs against the genome-wide background of multi-way contacts.

Here we develop Nanopore-HiChIP, a long-read H3K27ac chromatin conformation method that directly resolves high-order enhancer hubs in diploid genomes. By integrating allele-resolved chromatin architecture, transcription factor occupancy, bulk and single-cell transcriptomics, and transcriptional kinetic modelling, we show that naturally occurring genetic variation extensively remodels enhancer hubs but without generally producing allele-specific expression. Instead, enhancer hubs buffer regulatory perturbation through reciprocal compensation between transcriptional burst frequency and burst size, thereby preserving balanced transcriptional output despite substantial allelic differences in regulatory architecture. Finally, we show that this buffering architecture is preferentially deployed at haploinsufficient genes, providing a mechanistic explanation for how diploid genomes preserve allelic balanced dosage while tolerating pervasive regulatory variation.

### Nanopore-HiChIP resolves high-order enhancer hubs

To directly resolve high-order chromatin interactions at active regulatory elements, we adapted Pore-C by incorporating H3K27ac immunoprecipitation, developing Nanopore-HiChIP (Fig. 1A). Similar to Pore-C (*35*), Nanopore-HiChIP sequences intact proximity-ligation products using long-read nanopore sequencing, but selectively enriches for active chromatin interactions through H3K27ac immunoprecipitation. Since the protocol preserves intact concatemers without the sonication step used in conventional HiChIP, every ligation partner within a chromatin complex can be recovered simultaneously on a single sequencing read (Fig. 1A).

**Figure 1.**
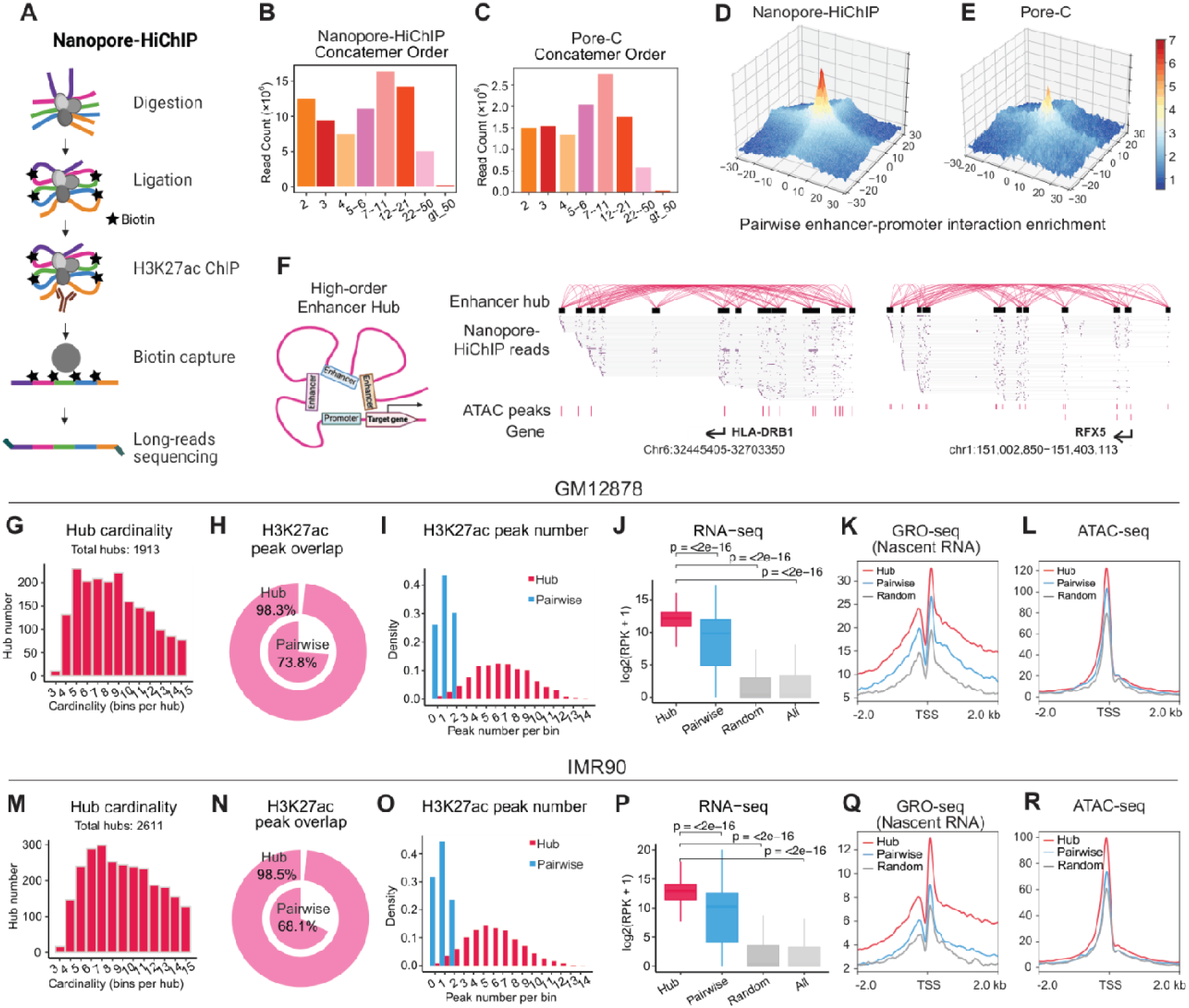
Nanopore-HiChIP resolves high-order enhancer hubs. **(A)** Nanopore-HiChIP workflow. Chromatin is digested and proximity-ligated in situ with biotin marking, followed by H3K27ac immunoprecipitation, biotin capture and long-read nanopore sequencing. Because ligation precedes immunoprecipitation and no sonication step is used, concatemers remain intact and every ligation partner within a chromatin complex is recovered on a single read. **(B, C)** Distribution of concatemer orders (number of ligation junctions per read) in Nanopore-HiChIP (B) and Pore-C (C) libraries from GM12878. Both assays recover a comparable range of high-order concatemers, indicating that adding an immunoprecipitation step does not compromise recovery of multi-way contacts. **(D, E)** Aggregate contact enrichment of Nanopore-HiChIP (D) and Pore-C (E) over enhancer–promoter pairs defined independently by conventional H3K27ac HiChIP. Axes show distance from the interaction anchors in 10 kb bins; colour indicates observed/expected contact enrichment. Nanopore-HiChIP shows stronger central enrichment, reflecting its selectivity for active regulatory chromatin. **(F)** Left, schematic of a high-order enhancer hub in which multiple enhancers converge simultaneously on a shared promoter. Right, two representative hubs at the HLA-DRB1 and RFX5 loci. Arcs show contacts among hub bins; each grey line below is a single Nanopore-HiChIP read, with purple marks indicating its constituent alignment fragments; ATAC-seq peaks and gene orientation are shown beneath. Individual reads span multiple hub bins, providing direct single-molecule evidence of simultaneous multi-enhancer engagement. **(G–L)** Characterization of the 1,913 enhancer hubs identified in GM12878. **(G)** Hub cardinality, the number of bins per hub. **(H)** Fraction of hub bins (outer ring) and pairwise enhancer anchors (inner ring) overlapping an H3K27ac ChIP-seq peak. **(I)** Number of H3K27ac peaks per bin for hub bins (red) and pairwise anchors (blue). Hub bins typically contain multiple peaks, whereas pairwise anchors contain none or one. **(J)** Expression of genes targeted by hubs, by pairwise enhancer contacts, and of random and genome-wide background gene sets. ****P < 0.0001, two-sided Wilcoxon rank-sum test is used versus Hub. **(K, L)** Metagene profiles of GRO-seq nascent transcription (K) and ATAC-seq accessibility (L) around the transcription start sites of the same gene classes. **(M–R)** The same analyses applied to the 2,611 enhancer hubs identified in IMR90, showing closely matching architectural and functional properties.

Nanopore-HiChIP libraries from GM12878 cells spanned a broad range of concatemer orders, with the largest fractions containing 7–11 and 12–21 ligation junctions (Fig. 1B). This distribution closely matched that obtained from Pore-C libraries prepared from the same cell line (Fig. 1C), indicating that inserting an immunoprecipitation step does not compromise recovery of high-order concatemers.

To assess whether Nanopore-HiChIP faithfully captures enhancer-promoter interactions, we aggregated Nanopore-HiChIP signal over enhancer–promoter pairs defined independently by conventional short-read H3K27ac HiChIP. Nanopore-HiChIP showed a sharp central enrichment that exceeded the enrichment observed in Pore-C over the same pairs (Fig. 1D,E), as expected given that Pore-C samples contacts genome-wide without selection for active chromatin. This confirms that long-read sequencing of intact concatemers does not sacrifice the pairwise information provided by short-read HiChIP while adding high-order structure.

Frequent multi-way co-occurrence alone does not inherently indicate cooperative chromatin organization. We therefore employed Chromunity (*35*) to identify statistically significant high-order assemblies by assessing whether observed multi-way contacts occur more frequently than anticipated based on their constituent pairwise interactions. We designate these statistically significant assemblies (FDR < 0.05) as high-order enhancer hubs. Notably, these hubs include the HLA, CD38, EBF1, CDK4, and IRF2 loci (*35*–*37*) (Fig. 1F, S1A), which have been previously identified as multi-way regulatory hubs through independent assays. At these established loci, Nanopore-HiChIP recovered both the corresponding genes and constituent enhancer bins, along with their co-occurrence patterns, thereby providing orthogonal validation for the identification of high-order enhancer hubs.

In total, we identified 1,913 high-order enhancer hubs in GM12878 cells (Table S1). These hubs contained between three and fifteen genomic bins, with most comprising six to nine bins (Fig. 1G), and spanned a median genomic distance of approximately 50 kb, with the majority extending less than 1 Mb (Fig. S1B). They were also enriched for active enhancer chromatin: over 95% overlapped with H3K27ac (Fig. 1H) and H3K4me1 (Fig. S1C) peaks, which is markedly higher than pairwise enhancer interactions. Individual hub bins of these hubs typically contained multiple H3K27ac (Fig. 1I) and H3K4me1 (Fig. S1D) peaks, whereas pairwise interactions most frequently contained only one or two. These observations indicate that high-order enhancer hubs are assembled from clusters of highly active enhancers rather than isolated enhancer elements.

Genes associated with high-order enhancer hubs were expressed at significantly higher levels than genes connected only through pairwise enhancer-promoter interactions, with both groups surpassing random and genome-wide backgrounds (Fig. 1J). We further examined nascent transcription and chromatin accessibility using GRO-seq, PRO-seq and ATAC-seq. Promoters associated with hubs displayed stronger promoter-proximal GRO-seq (Fig. 1K), PRO-seq (Fig. S1E) and ATAC-seq (Fig. 1L) signal than those regulated by pairwise interactions, indicating that the elevated expression originates from increased transcriptional activity rather than post-transcriptional regulation. Together, these observations establish high-order enhancer hubs as highly active multi-enhancer regulatory assemblies associated with elevated transcriptional output.

Subsequently, we applied the same analysis pipeline to IMR90 fibroblasts, identifying 2,611 high-order enhancer hubs (Table S1) that exhibited architectural properties closely resembling those observed in GM12878, including hub cardinality and genomic span (Fig. 1M, Fig. S1F). The hub bins displayed substantially greater H3K27ac (Fig. 1N,O) and H3K4me1 (Fig. S1G,H) occupancy than pairwise enhancer interactions. Consistently, genes associated with high-order enhancer hubs showed significantly higher expression than genes linked only through pairwise enhancer interactions (Fig. 1P), which was accompanied by elevated promoter-proximal GRO-seq, PRO-seq and ATAC-seq signal (Fig. 1Q,R, Fig. S1I). These findings indicate that the defining structural and functional properties of enhancer hubs are reproducibly conserved across distinct cell types, establishing enhancer hubs as a general organizational framework for active gene regulation and laying the foundation for understanding their biological functions.

### Enhancer hubs undergo widespread allele-specific remodeling

Having identified high-order enhancer hubs, we next asked whether the two homologous copies of a hub within the same nucleus are structurally equivalent. Long-read concatemers make this question directly addressable, as individual reads spanning one or multiple heterozygous variants can be assigned unambiguously to a parental haplotype. We therefore phased Nanopore-HiChIP concatemers using the ENCODE reference haplotypes for each cell line and compared enhancer hubs composition changes between homologous chromosomes.

Allelic hub differences are not a binary presence-or-absence property, since contact frequency varies continuously and is measured with sampling noise. We first compared the overall contact strength of each hub between homologous chromosomes. This global comparison identified few significant allelic differences, suggesting that natural genetic variation does not generally induce a uniform weakening of an entire hub.

Given that a hub comprises multiple constituent bins, we investigated whether individual bins contributed differently to the overall contact profile of the same hub between homologous chromosomes. To test this, we adapted a differential usage framework (DEXSeq (*38*)), treating each hub as a unit and its constituent bins as distinct features. This approach accounts for baseline differences in total hub strength, allowing us to identify "lost" bins whose proportional contribution to the hub was significantly reduced on one allele (Fig. 2A). Within individual hubs, lost bins demonstrated significantly greater allelic contact differences than other retained bins (Fig. 2B, 2C). Hubs with lost bins predominantly on allele 2 were classified as allele 1-biased, whereas those with lost bins predominantly on allele 1 were classified as allele 2-biased; hubs with losses on both alleles were classified as mixed (Fig. 2B, 2C). Representative loci illustrate the underlying single-molecule evidence, with haplotype-specific concatemers consistently including or excluding individual regulatory elements that contain heterozygous sequence variants (Fig. 2D, E, S2).

**Figure 2.**
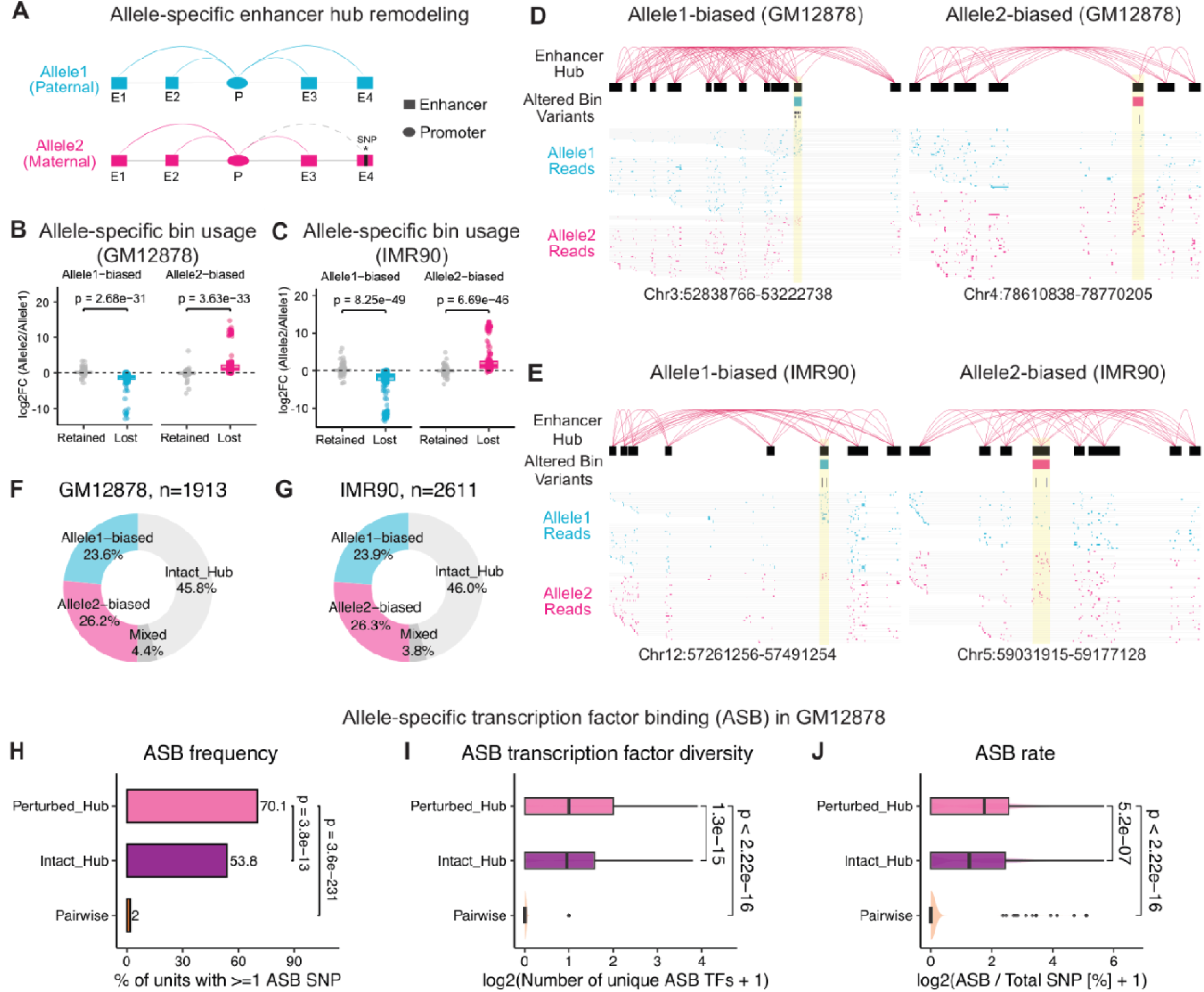
Enhancer hubs undergo widespread allele-specific remodeling. **(A)** Schematic of allele-specific enhancer hub remodeling. A heterozygous variant within one enhancer disrupts its participation in the hub on the maternal allele (dashed arc), while the same element remains engaged on the paternal allele. **(B, C)** Allelic contact change at hub bins in GM12878 (B) and IMR90 (C). Each dot is one bin belonging to a perturbed hub. The y axis gives the log ratio of that bin’s contact frequency between alleles (allele 2 / allele 1): zero means the bin contributes equally on both alleles, positive values mean it contributes more on allele 2, and negative values more on allele 1. Along the x axis, bins are grouped by whether DEXSeq identified them as differentially used between alleles ("Lost", coloured) or not ("Retained", grey). The two facets separate hubs classified as allele 1–biased from those classified as allele 2–biased. Within each perturbed hub, bins identified by DEXSeq as differentially used ("Lost") are compared with bins retained on both alleles ("Retained") from the same hub. Two-sided Wilcoxon rank-sum test is used. **(D, E)** Representative allele-biased hubs in GM12878 (D) and IMR90 (E). Arcs show hub contacts; the altered bin is highlighted; individual reads assigned to allele 1 (blue) and allele 2 (red) are shown below. Reads from one haplotype span the altered bin while reads from the other omit it. **(F, G)** Hub classification in GM12878 (F) and IMR90 (G). Hubs were designated as Intact if no bins were lost, whereas those with at least one lost bin were classified as Perturbed. Within the perturbed category, hubs were further categorized as Allele1-biased or Allele2-biased if all lost bins shifted towards the same allele, or as Mixed if the bins shifted in both directions. **(H–J)** Allele-specific transcription factor binding (ASB) in GM12878, comparing perturbed hubs, intact hubs and pairwise enhancer–promoter interactions. **(H)** Percentage of units containing at least one ASB variant. **(I)** Number of distinct transcription factors showing ASB per unit. **(J)** ASB rate, the fraction of phased heterozygous variants within each unit called as ASB. Using heterozygous sites as the denominator controls for differences in sequencing coverage between region classes. Two-sided Wilcoxon rank-sum test is used.

Allelic hub perturbation is notably prevalent. Of the 1,913 enhancer hubs identified in GM12878, 54.2% were categorized as perturbed hubs (Table S2). This includes 23.6% biased towards allele 1, 26.2% towards allele 2, and 4.4% exhibiting perturbations in both directions (Fig. 2F). A similar distribution of perturbed and intact hubs was observed in IMR90 (Fig. 2G; Table S2). By comparing bin usage strictly within the same hub, we controlled for the broader genomic locus, target gene, and local chromatin environment. This approach allowed us to directly map allelic structural divergence to the specific bins driving the perturbation. Collectively, these findings indicate that allelic enhancer hub differences are driven by focal remodeling of distinct regulatory elements.

To determine that these differences reflect genuine sequence-directed regulatory changes rather than technical variation in contact detection, we asked whether they coincide with allelic imbalance in transcription factor occupancy. We intersected enhancer hubs with allele-specific transcription factor binding (ASB) events from ADASTRA (*39*), using phased heterozygous variants as the denominator to mitigate potential bias due to differential sequencing coverage across regions. Perturbed hubs were significantly more enriched for ASB compared to both intact hubs and pairwise enhancer interactions. 70.1% of perturbed hubs contained at least one ASB event, compared with 53.8% of intact hubs and only 2% of pairwise interactions (Fig. 2H). Additionally, perturbed hubs also engaged a broader repertoire of transcription factors (Fig. 2I), and a significantly larger fraction of heterozygous variants within these hubs exhibited allele-specific binding (Fig. 2J). Taken together, these results show that heterozygous genetic variation extensively remodels enhancer hubs between homologous chromosomes, characterized by focal, allele-specific changes in constituent regulatory elements and transcription factor occupancy.

### Allele-specific enhancer hubs rarely produce allele-specific expression

Having established that heterozygous variants extensively remodel enhancer hubs between homologous chromosomes (Fig. 2), we next asked whether these regulatory perturbations propagate to transcriptional output. We therefore quantified allele-specific expression (ASE) using bulk RNA-seq, identifying 715 significant ASE genes in GM12878 and 807 in IMR90 (Fig. 3A,B). The analysis recovered numerous well-established imprinted genes, including KCNQ1OT1, SNRPN, PEG10 and MEG8 (*40*, *41*), confirming the robust detection of genuine allelic imbalance.

**Figure 3.**
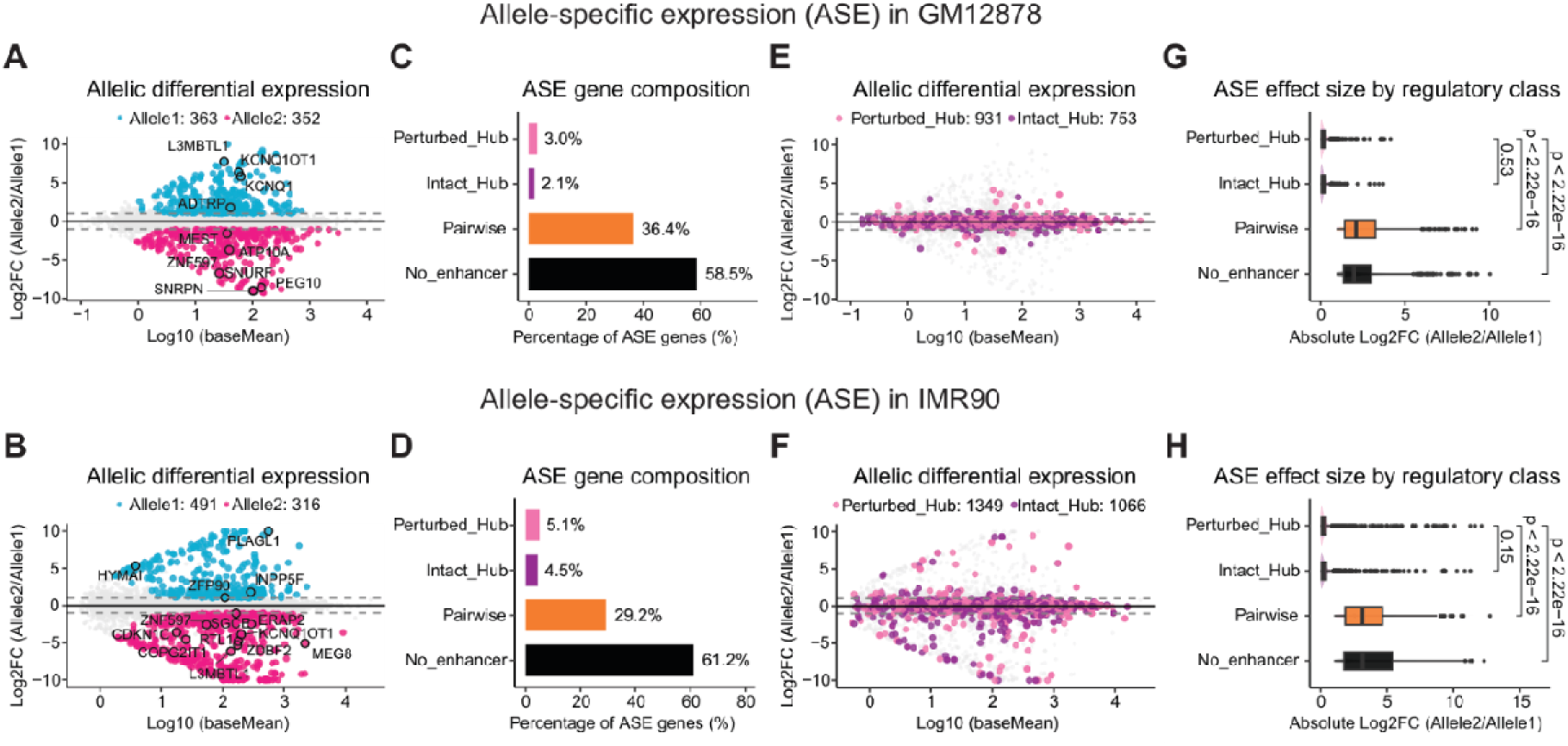
Allele-specific enhancer hubs rarely produce allele-specific expression. **(A, B)** Allelic differential expression from bulk RNA-seq in GM12878 (A) and IMR90 (B). Coloured points are genes with significant allele-specific expression (ASE); dashed lines mark two-fold allelic difference. Labelled genes are known imprinted loci recovered as expected, confirming that the analysis detects genuine allelic imbalance. **(C, D)** Composition of the ASE gene set by regulatory class in GM12878 (C) and IMR90 (D). Hub-target genes account for a small minority of ASE genes. Within-class ASE rates, which control for the different sizes of the four classes, are shown in Fig. S3A, B. **(E, F)** Allelic differential expression restricted to hub-target genes in GM12878 (E) and IMR90 (F), plotted over the genome-wide background (grey). Coloured points are genes associated with perturbed (pink) and intact (purple) hubs. Despite extensive allelic hub remodeling in perturbed hubs, these genes cluster around balanced expression across the full expression range. **(G, H)** Absolute allelic fold change by regulatory class in GM12878 (G) and IMR90 (H). Perturbed and intact hub genes show comparably small allelic differences, both substantially smaller than pairwise-regulated and enhancer-free genes. Two-sided Wilcoxon rank-sum test is used.

Contrary to the conventional expectation that cis-regulatory perturbations influence gene expression, perturbed hub-target genes were notably absent from the ASE set. Genes associated with both perturbed and intact enhancer hubs constituted only 4.1% of ASE genes in GM12878 and 9.6% in IMR90 (Fig. 3C,D). The majority of ASE genes were linked to pairwise enhancer-promoter interactions or lacked detectable enhancer contacts (Fig. 3C,D). Given the substantial differences in gene numbers across these four regulatory classes, we calculated the ASE rate within each class. The results were consistent: only 2.4% of perturbed-hub genes and 2.1% of intact-hub genes were classified as ASE in GM12878, compared to 9.7% of pairwise-regulated genes and 6.6% of enhancer-free genes, with a similar pattern observed in IMR90 (Fig. S3A,B). Therefore, genes regulated by enhancer hubs exhibit markedly lower rates of allele-specific expression than genes without hub regulation.

The magnitude of allelic expression differences exhibited a consistent pattern. In allele-specific MA plots, both perturbed and intact hub genes clustered tightly around balanced expression across the entire dynamic range in GM12878 (Fig. 3E), with the majority showing less than a two-fold allelic difference; IMR90 demonstrated a similar distribution (Fig. 3F). When quantified across regulatory classes, perturbed and intact hub genes displayed indistinguishable allelic fold changes, both significantly smaller than those observed for pairwise-regulated and enhancer-free genes (Fig. 3G, 3H). These results strongly indicate that the expression is preserved even where hub structure is demonstrably altered.

The depletion of ASE at hub-target genes cannot be attributed to limited statistical power. Hub-target genes showed significantly greater allelic read depth than pairwise-regulated or enhancer-free genes in both cell types (Fig. S3C,D), along with comparable or greater numbers of heterozygous exonic variants (Fig. S3E,F), the two primary determinants of statistical power for detecting allele-specific expression. Technically, hub-target genes should have been the easiest class in which to detect ASE, yet they consistently yielded the fewest ASE events.

Together, these analyses reveal the central paradox addressed by this study. In contrast to the conventional view that genetic variants that disrupt transcription factor binding or local chromatin structure are effectively transmitted downstream, leading to allele-specific gene expression. Genetic variation targeting enhancer hubs can extensively remodel high-order chromatin architecture and transcription factor occupancy without resulting in corresponding expression differences. The genetic perturbation therefore reaches the regulatory layer, but does not reach transcriptional output. Thus, high-order enhancer hubs act as a buffering architecture, absorbing the consequences of allelic genetic variation to maintain stable and balanced gene expression.

### Enhancer hubs buffer allelic regulatory perturbation through transcriptional kinetic compensation

Having established the buffering capacity of enhancer hubs, we aimed to elucidate the specific mechanisms within the transcription process where these genetic perturbations are absorbed. Given that bulk RNA-seq only measures aggregate expression across cells, we employed allele-resolved single-cell RNA-seq to resolve cell-to-cell transcription variation from each allele. Consistent with bulk measurements, merged single-cell data indicated that allelic differences in aggregate expression remained centered on zero for perturbed-hub genes and were indistinguishable from intact-hub genes. In contrast, pairwise-regulated and enhancer-free genes exhibited substantially larger allelic differences in both cell types (Fig. S4A, S4B). Despite nearly identical aggregate expression, the underlying single-cell transcriptional distributions were markedly different. At representative gene loci associated with perturbed hubs, one allele was transcriptionally silent in more cells but produced many transcripts when active, whereas the other allele was active in a larger fraction of cells but generated fewer transcripts per activation event (Fig. 4A, S4C). This pattern was not limited to individual examples. Across perturbed-hub genes, the two alleles systematically differed in both the fraction of transcriptionally silent cells and the number of transcripts produced per expressing cell (Fig. S4D).

**Figure 4.**
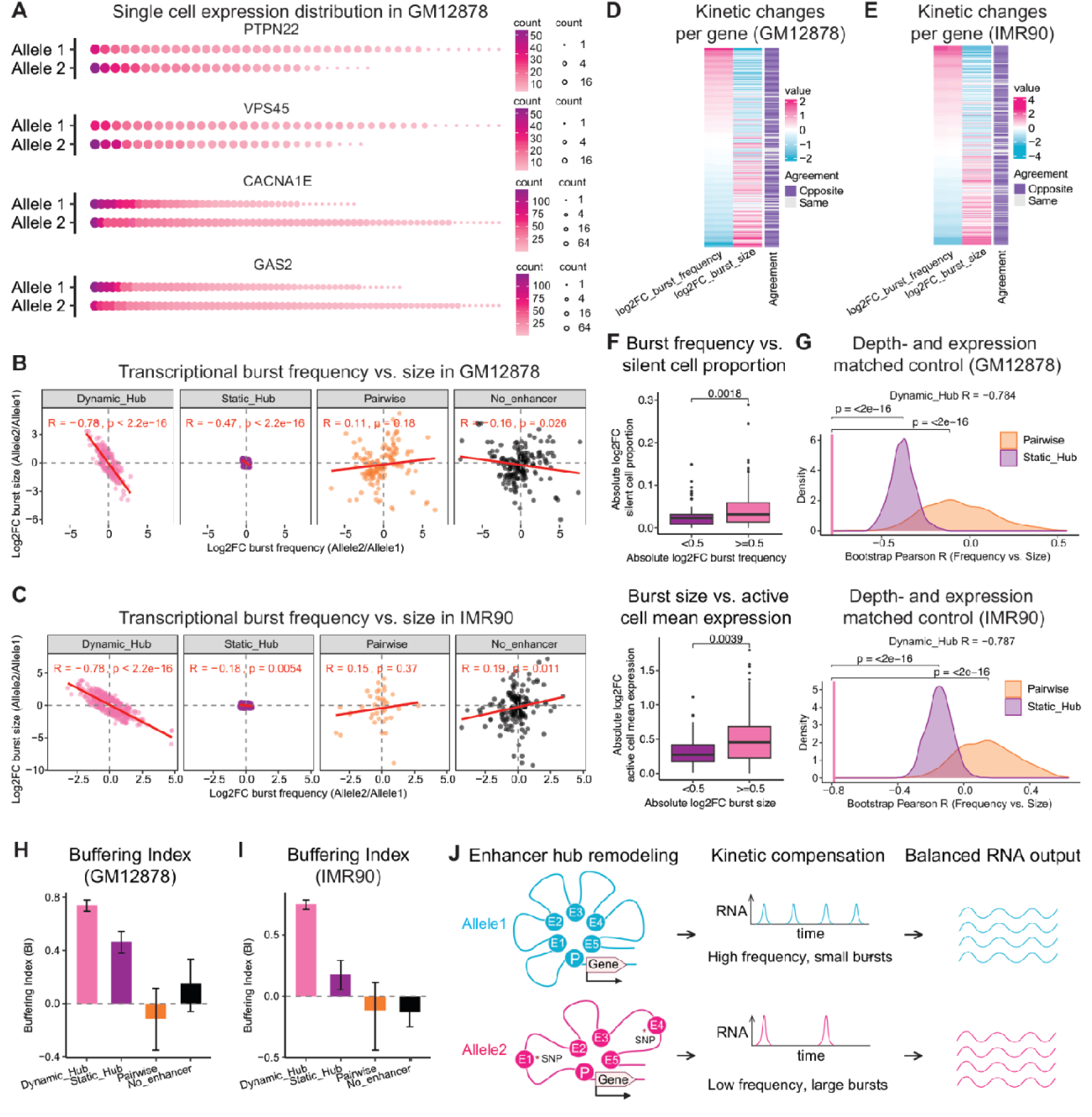
Enhancer hubs buffer allelic regulatory perturbation through transcriptional kinetic compensation. **(A)** Allele-resolved single-cell expression distributions for four representative hub-target genes in GM12878. Each dot is one cell, ordered by transcript count; dot size and colour indicate count. For the same gene, one allele is silent in more cells but produces more transcripts where active, while the other is active in more cells with fewer transcripts each. **(B, C)** Allelic change in burst frequency plotted against allelic change in burst size, in GM12878 **(B)** and IMR90 **(C)**. Each dot is one gene; both axes are log ratios between alleles (allele 2 / allele 1). Panels separate the four regulatory classes: dynamic hubs are perturbed-hub genes whose kinetics differ between alleles (|log FC| > 0.5 in either parameter), static hubs are perturbed-hub genes below that threshold, and pairwise-regulated and enhancer-free genes serve as comparison classes. The red line is a linear fit; R and P are Pearson correlation. A negative slope means that the allele firing less often produces more transcripts per burst. **(D, E)** The same allelic kinetic changes shown gene by gene for dynamic hubs, in GM12878 **(D)** and IMR90 **(E)**. Each row is one gene, sorted by its change in burst frequency. The two columns give the allelic log change in burst frequency and in burst size, coloured pink for positive and blue for negative (scale at right). The narrow bar on the far right indicates, for each gene, whether the two parameters changed in opposite directions (purple, "Opposite") or the same direction (lightgrey, "Same"). **(F)** The boxplot illustrates that the fitted kinetic parameters accurately represent the raw single-cell data for GM12878. Genes are categorized based on the magnitude of their fitted allelic kinetic change (x-axis: |log FC| below or at/above 0.5) and are evaluated using a raw distributional measure derived directly from counts without model fitting (y-axis). In the top panel, genes exhibiting larger fitted changes in burst frequency demonstrate greater allelic differences in the proportion of cells where the gene is inactive. In the bottom panel, genes with more substantial fitted changes in burst size display greater allelic differences in the mean transcript count among cells that express the gene. Two-sided Wilcoxon rank-sum test is used. **(G)** Pairwise-regulated genes (orange) and static hubs (purple) were matched to dynamic hubs on expression level and allelic read depth, and the frequency–size correlation recomputed on 1,000 bootstrap resamples of each matched set; the curves are the resulting distributions of Pearson R. The vertical pink line marks the correlation actually observed for dynamic hubs, which falls outside both distributions. **(H, I)** Buffering Index for each regulatory class in GM12878 **(H)** and IMR90 **(I).** The index is the proportion of total kinetic variance cancelled by reciprocal covariance between burst frequency and burst size, BI = –2 Cov / (Var_f + Var_b). A value of 0 means the two parameters vary independently and their changes pass through to expression; a value of 1 means they cancel exactly and expression is unchanged. Bars are coloured by class as in (B) and (D); error bars are 95% bootstrap confidence intervals, and an interval crossing zero indicates no detectable buffering. **(L)** Model. At enhancer hubs, the same class of perturbation is absorbed by reciprocal adjustment of burst frequency and burst size, so the perturbation reaches the regulatory layer but not transcriptional output.

These two features of the single-cell distribution are empirical signatures of transcriptional bursting dynamics: the fraction of silent cells reflecting how often a gene fires (burst frequency), and the transcript count in active cells reflecting how much it produces per firing (burst size). We therefore inferred allele-specific burst parameters under the classical two-state model of gene expression (*42*). Among perturbed-hub genes exhibiting substantial kinetic changes (|log FC| > 0.5 in burst frequency or burst size; hereafter referred to as dynamic hubs), burst frequency and burst size changed in a strikingly reciprocal manner. Alleles firing less frequently consistently produced more transcripts per burst, whereas alleles firing more frequently generated proportionally smaller bursts (R = –0.78, *P* < 2.2 × 10 ¹; Fig. 4B). This pattern was unique to perturbed hubs. Neither pairwise-regulated genes (R = 0.11) nor enhancer-free genes (R = – 0.16) exhibited evidence of reciprocal compensation despite displaying comparable kinetic perturbations. Consequently, transcriptional kinetic changes in these genes propagated directly to expression differences, explaining why they constitute the majority of allele-specific expression events (Fig. 3G). The same pattern was independently reproduced in IMR90, where dynamic hubs displayed an equally strong inverse relationship (R = –0.78), whereas both comparison groups again lacked evidence of compensation (Fig. 4C). This reciprocal relationship is also evident across individual genes ordered by burst-frequency difference in both cell types (Fig. 4D,E).

Importantly, this reciprocal relationship was not attributable to technical reasons. First, the inferred parameters faithfully recapitulated the underlying single-cell distributions: genes exhibiting larger allelic differences in burst frequency showed correspondingly larger shifts in silent-cell fraction, whereas genes with larger burst-size differences displayed larger changes in transcript output per expressing cell (Fig. 4F, S4E). Hence, the inferred kinetic parameters capture genuine features of the observed single-cell distributions rather than representing artefacts of model fitting. Second, matching pairwise-regulated genes and static hubs to dynamic hubs for both expression level and allelic sequencing depth had little effect on the observed compensation, which remained well outside the bootstrapped null distributions of both matched control groups in GM12878 and IMR90 (Fig. 4G). Furthermore, although static hubs exhibited a weak negative correlation, the associated kinetic variation was nearly an order of magnitude smaller than that observed in dynamic hubs, indicating that this correlation is dominated by the joint estimation uncertainty of the two parameters rather than by biological variation. Together, these analyses confirm that reciprocal changes in burst frequency and burst size of dynamic hubs represent a genuine biological property of enhancer hubs, rather than a consequence of statistical inference.

To quantify the extent to which transcriptional kinetic perturbations are prevented from propagating to expression, we decomposed the variance of allelic mean expression into the contributions from burst frequency, burst size and their covariance, defining a Buffering Index (BI) as the fraction of total kinetic variance cancelled by reciprocal covariance between the two parameters (BI = –2Cov/(Var_f + Var_b)). Dynamic hubs buffered the majority of transcriptional kinetic variation (BI = 0.74; Fig. 4H). In contrast, neither pairwise-regulated genes nor enhancer-free genes showed evidence of significant buffering, with confidence intervals overlapping zero.

IMR90 replicated the same ordering, with dynamic hubs achieving an even higher buffering index (BI = 0.75; Fig. 4I). Thus, the majority of the transcriptional kinetic perturbation introduced by naturally occurring allelic regulatory variation is absorbed within the transcription cycle before it can propagate to steady-state gene expression.

These results resolve the paradox revealed in Fig. 2 and Fig. 3. Although allelic sequence variation rewires enhancer hubs and alters transcription factor occupancy, it does not directly translate into expression divergence. Instead, hub-target genes absorb these perturbations through reciprocal changes in burst frequency and burst size, preserving overall transcriptional output despite substantial changes in transcriptional kinetics. Kinetic compensation therefore provides the mechanistic basis by which enhancer hubs buffer the transcriptional consequences of allele-specific regulatory perturbation (Fig. 4J).

### Enhancer hubs safeguard haploinsufficient genes from regulatory perturbation

Having established that enhancer hubs buffer allele-specific regulatory perturbations through kinetic compensation, we next asked why such a buffering system is required and which genes it serves. To address this question, we compared enhancer hubs between GM12878 and IMR90 cells. Hub architecture proved to be highly cell-type-specific: only 276 hubs were shared between GM12878 and IMR90, representing just 7.3% of the union (Fig. 5A). The genes targeted by these hubs reflected the biological identity of each cell type. GM12878 hub genes were enriched for B-cell receptor signalling, antigen receptor signalling and NF-κB pathways, whereas IMR90 hub genes were enriched for receptor tyrosine kinase signalling, cell migration and endothelial migration (Fig. 5B). Disease ontology analysis revealed the same cell identity specificity, linking GM12878 hubs predominantly to leukemias and lymphomas, whereas IMR90 hubs were preferentially associated with solid tumors, as well as tumor progression and metastasis (Fig. S5A). These observations indicate that enhancer hubs are not constitutive architectural features of the genome, but are assembled in a cell-type-specific manner to regulate the transcriptional programs that define cellular identity.

**Figure 5.**
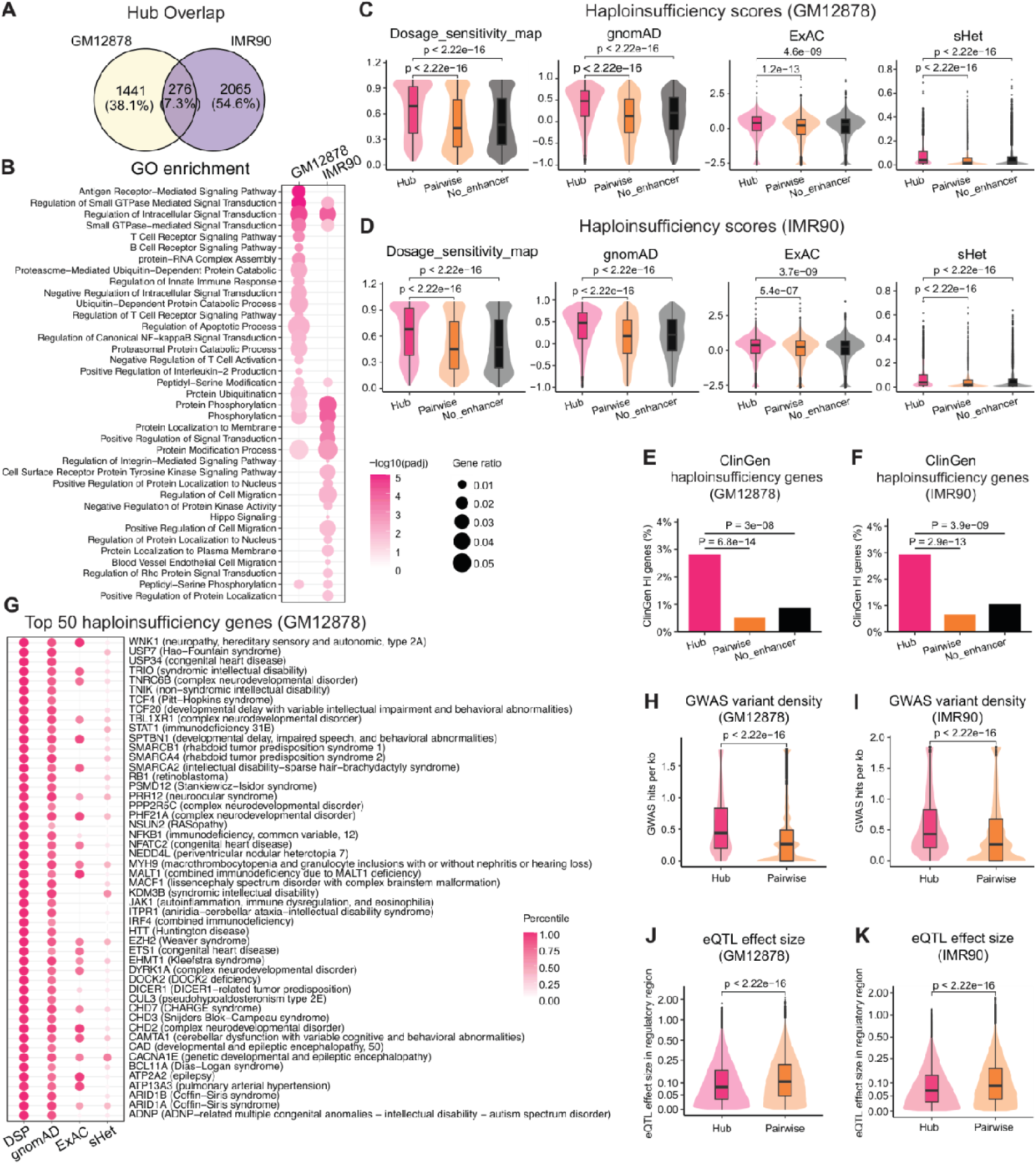
Enhancer hubs safeguard haploinsufficient genes from regulatory perturbation. **(A)** Overlap between the enhancer hubs identified in GM12878 (yellow) and IMR90 (purple), with counts and the percentage of the union in each sector. **(B)** Gene Ontology biological process terms enriched among hub-target genes, shown separately for the two cell lines. Dot size is the gene ratio, the proportion of genes in that term found among hub targets; colour is enrichment significance as –log (adjusted P), with darker pink indicating stronger enrichment. A missing dot means the term was not enriched in that cell line. **(C, D)** Haploinsufficiency scores for genes in each regulatory class, in GM12878 **(C)** and IMR90 **(D)**. The four sub-panels are independently derived measures and are on different scales: the dosage sensitivity map (haploinsufficiency score) and gnomAD loss-of-function intolerance probability both give probabilities from 0 to 1; the ExAC loss-of-function z score is a signed deviation statistic; and sHet estimates the selection coefficient against heterozygous loss of function. In all four, higher values indicate greater intolerance to losing one copy. Violins show the full distribution with boxplots inside; hub-target genes are pink, pairwise-regulated genes orange, enhancer-free genes grey. Two-sided Wilcoxon rank-sum test is used. **(E, F)** Percentage of genes in each class that appear on the ClinGen list of experimentally validated haploinsufficient genes, in GM12878 **(E)** and IMR90 **(F)**. Unlike the scores in (C, D), which are inferred from population sequencing, ClinGen entries are curated from clinical evidence. Fisher’s exact test. **(G)** The 50 hub-target genes in GM12878 with the highest haploinsufficiency scores from the dosage sensitivity map, with the disease caused by their loss of function given in parentheses. Each row is a gene and each column one of the four measures from (C); dot colour gives that gene’s percentile rank within the measure, from white (low) to dark pink (high). **(H, I)** The density of GWAS Catalog associations within regulatory regions is presented for GM12878 (H) and IMR90 (I). The counts are normalized per kilobase of regulatory sequence due to the varying sizes of hub regions and pairwise anchors. The violin plots illustrate the distribution across regions, with boxplots embedded within; hub regions are depicted in pink, while pairwise enhancer anchors are shown in orange. Two-sided Wilcoxon rank-sum test is used. **(J, K)** The absolute effect size of cis-eQTLs located within identical regulatory regions is analyzed using tissue-matched GTEx data for GM12878 (J) and IMR90 (K). Each observation represents an eQTL variant, categorized into hub or pairwise regions based on genomic position. The y-axis denotes the effect size, which is the magnitude of the variant’s effect on the expression of its target gene, presented on a square-root scale. The color scheme corresponds to that in (H, I). Two-sided Wilcoxon rank-sum test is used.

Genes specifying cell identity depends critically on gene dosage rather than mere presence. For many of these genes, expression from a single allele is insufficient to maintain cellular function and cause disease (*1*–*6*). We therefore asked whether enhancer hubs preferentially deployed to regulate haploinsufficient genes. Across four independent measures of haploinsufficiency— including the dosage sensitivity map (haploinsufficiency index) (*1*), the gnomAD loss-of-function intolerance probability (*6*), the ExAC loss-of-function z score (*43*), and sHet, an estimate of the selection coefficient against heterozygous loss of function (*44*) (Table S3)—hub target genes consistently scored significantly higher than both pairwise-regulated genes and genes lacking enhancer regulation in GM12878 (Fig. 5C). The same pattern was independently reproduced in IMR90 (Fig. 5D). Hub targets were also significantly enriched among experimentally curated ClinGen (*45*) haploinsufficient genes in both cell types (Fig. 5E,F; Table S3). Despite their distinct derivations and statistical assumptions, all four dosage-sensitivity metrics converged on the same conclusion: enhancer hubs are preferentially deployed on genes whose balanced allelic dosage must be tightly maintained.

The identity of these genes further reinforces this interpretation. The most haploinsufficient hub targets in GM12878 are chromatin regulators and developmental transcription factors underlying well-characterised haploinsufficient syndromes, including CHD7, ARID1A, ARID1B, EHMT1, TCF4, SMARCB1, SMARCA4, EZH2 and RB1, together with lineage-specific immune regulators such as STAT1, NFKB1, JAK1 and IRF4 in GM12878 (Fig. 5G). Similar enrichment was observed in IMR90, where hub targets included CREBBP, EP300, SHANK3, KAT6A and TRIO, alongside connective-tissue genes such as FBN1, COL4A1 and COL5A1 (Fig. S5B). For genes of this class, monoallelic expression frequently results in clinically recognisable phenotypes, highlighting the biological importance of maintaining proper allelic dosage from both parental alleles.

To better understand why enhancer hubs evolved to safeguard these haploinsufficient genes, we further examined the mutational burden across hub architecture. Hub regulatory regions contained significantly higher densities of GWAS-associated variants (*46*) (Fig. 5H,I) and ClinVar pathogenic variants (*47*) (Fig. S5C,D) than pairwise enhancer anchors in GM12878, indicating hub regions accumulate greater burden of disease-associated regulatory variation. Surprisingly, despite carrying substantially more regulatory variation, variants located within hub regions exhibited significantly smaller cis-eQTL effect sizes than variants located within pairwise enhancers in matched GTEx tissues (*12*, *48*) (Fig. 5J,K), indicating that the buffering we measured allele by allele in single cells is detectable as an attenuation of regulatory effect across human populations. These apparently opposing observations suggest that enhancer hubs may have evolved as an architectural safeguard for preserving biallelic transcriptional robustness, allowing haploinsufficient genes to endure pervasive cis-regulatory variation without compromising balanced expression from two genetically distinct parental alleles.

## Discussion

A central paradox of diploid gene regulation is the extensive divergence in cis-regulatory landscapes between the two parental chromosomes, which nonetheless results in similar transcriptional outputs (*12*–*17*). Our study provides a mechanistic explanation for this paradox by identifying high-order enhancer hubs as a regulatory architecture that buffers naturally occurring cis-regulatory variation before it propagates to gene expression. We show that genetic variation can remodel enhancer-hub architecture and alter transcription factor occupancy between homologous chromosomes, yet these regulatory perturbations are largely prevented from causing allelic differences in steady-state expression. Instead, the perturbations are absorbed through reciprocal changes in transcriptional burst frequency and burst size, allowing transcriptional output to remain balanced between parental alleles. Thus, enhancer hubs do not eliminate the regulatory consequences of genetic variation; rather, they accommodate changes in three-dimensional regulatory organization and transcription factor occupancy through compensatory changes in transcriptional kinetics.

This discovery advances the current understanding of enhancer-redundancy-mediated buffering. Previous research has shown that redundant enhancer architectures can buffer genetic perturbations, as mean gene expression often remains largely unchanged despite the loss or weakening of individual regulatory elements(*22*–*25*). However, our findings indicate that enhancer-hub-mediated buffering is not simply preserving a static expression state; rather, it is an active redistribution of transcriptional dynamics that preserves overall transcriptional output. At allelically perturbed hubs, changes in burst frequency are counterbalanced by reciprocal changes in burst size, ensuring that perturbations in one kinetic dimension are absorbed by the other before affecting steady-state transcript abundance. This insight redefines the concept of regulatory robustness: stable expression does not imply stable regulation. Rather, transcriptional output can remain robust precisely because the underlying kinetic state is allowed to change. By revealing kinetic compensation as a mechanistic basis of enhancer-hub buffering, our findings provide a dynamic framework for understanding how high-order regulatory architectures absorb cis-regulatory perturbations.

The preferential association of enhancer hubs with haploinsufficient genes provides a biological context for why such a buffering system is important. Haploinsufficient genes are highly sensitive to changes in allelic dosage, yet their regulatory regions are also enriched for naturally occurring regulatory variation, including common disease-associated and pathogenic variants. This creates a fundamental evolutionary tension: regulatory sequences must remain sufficiently flexible to evolve, while the expression of haploinsufficient genes must remain sufficiently stable to preserve function. Our findings suggest that enhancer hubs provide a solution to this tension. By buffering the transcriptional consequences of cis-regulatory variation, enhancer hubs decouple sequence evolution from transcriptional stability, allowing regulatory sequences to diversify while safeguarding the expression of genes where dosage balance is essential. This principle may extend more broadly to dosage-sensitive genes whose function depends critically on maintaining appropriate expression levels. In this context, enhancer hubs may represent an evolutionary strategy that allows regulatory diversity and functional constraint to coexist within the same genome.

Future studies should focus on how enhancer hubs implement and delimit kinetic compensation. Firstly, how are changes in allele-specific enhancer architecture translated into reciprocal changes in burst frequency and burst size? One hypothesis is that allelic remodeling of enhancer hubs alters the local recruitment or redistribution of transcription factors and coactivators, including complexes involved in enhancer–promoter communication. This, in turn, could alter the organization or dynamics of transcriptional condensates or other high-order regulatory assemblies. Understanding how these molecular changes propagate from enhancer occupancy to transcriptional dynamics will require direct measurements of coactivator organization and transcription in real time. Secondly, previous research suggests that individual elements within an enhancer hub are not functionally equivalent (*49*, *50*). Different elements may occupy distinct functional roles within a hub, with some contributing primarily to transcriptional output, others providing buffering capacity, and still others facilitating communication among regulatory elements. Defining this functional hierarchy will be important for understanding why some perturbations remain within the buffering capacity of a hub whereas others escape it. These questions point toward a broader view of enhancer hubs not as collections of interchangeable enhancers, but as dynamic regulatory architectures whose internal organization determines how genetic variation is translated, or not translated, into transcriptional change.

## Materials and Methods

### 1. Experimental procedures

#### Cell culture

GM12878 and IMR90 cells were cultured following ENCODE consortium guidelines. GM12878, a lymphoblastoid line, was grown in suspension in RPMI 1640 with 15% fetal bovine serum, 2 mM L-glutamine and penicillin/streptomycin at 37°C and 5% CO₂, and passaged at 3–8 × 10 cells/mL. IMR90, a primary human fetal lung fibroblast line, was grown adherently in DMEM with 10% fetal bovine serum, 2 mM L-glutamine and penicillin/streptomycin under the same conditions, passaged at 70–80% confluency with trypsin-EDTA, and used within the ENCODE-recommended passage range to avoid replicative senescence. Both lines were confirmed mycoplasma-free before use.

#### Nanopore-HiChIP library preparation and sequencing

Nanopore-HiChIP was performed by adapting the Oxford Nanopore Technologies (ONT) Pore-C protocol, version 5 (2 March 2022), to incorporate immunoprecipitation of H3K27ac-associated chromatin following proximity ligation while preserving long proximity-ligated DNA molecules for nanopore sequencing. Approximately 1–5 × 10^6^ cells in suspension were used as input for each experiment. Unless otherwise specified below, Pore-C steps were performed according to the ONT Pore-C v5 protocol.

##### Cross-linking and nuclei isolation

Approximately 1–5 × 10^6^ cells were harvested and cross-linked in 1% formaldehyde (Fisher Scientific, cat. #BP531-25) for 10 min at room temperature. Cross-linking was quenched with 125 mM glycine (MilliporeSigma, cat. #50046-50G), and cells were washed with ice-cold PBS (Gibco, cat. #14190-136).

Cells were permeabilized according to the ONT Pore-C v5 protocol. The permeabilization buffer was supplemented with 3 mM MgCl_2_ (MilliporeSigma, cat. #M4880-100G), 5 mM sodium butyrate (MilliporeSigma, cat. #B5887-1G), and EDTA-free protease inhibitor cocktail (MilliporeSigma, cat. #11873580001). Cells were incubated on ice for the permeabilization period specified in the protocol. Subsequent enzymatic reactions were performed on intact, cross-linked nuclei to preserve chromatin contacts.

##### In situ restriction digestion

Nuclei were permeabilized with a brief SDS (Invitrogen, cat. #15553027) treatment (0.1% final) and quenched with Triton X-100 (MilliporeSigma, cat. #T8787) to a final concentration of 1%.

Chromatin was digested in situ with the four-cutter restriction endonuclease NlaIII (NEB, cat. #R0125) according to the Pore-C protocol. The restriction digest was additionally supplemented with 5 mM sodium butyrate and EDTA-free protease inhibitor cocktail to preserve histone acetylation and minimize proteolysis during the extended digestion.

##### Proximity ligation

Following restriction digestion, proximity ligation was performed according to the ONT Pore-C v5 protocol using T4 DNA ligase and the corresponding ligase buffer. Sodium butyrate (5 mM) and EDTA-free protease inhibitor cocktail were maintained during the ligation reaction.

This in situ ligation step joins restriction fragments held in spatial proximity within individual cross-linked nuclei, generating proximity-ligated molecules containing multiple chromatin contacts that can subsequently be resolved by long-read nanopore sequencing.

Following proximity ligation, nuclei were pelleted at 500 × g for 10 min and the ligation supernatant was removed.

##### H3K27ac chromatin immunoprecipitation

Nuclei were resuspended in ChIP lysis buffer containing 20 mM Tris-HCl (pH 8.0), 150 mM NaCl, 1% Triton X-100, 0.05% SDS, 0.1% sodium deoxycholate, 1 mM EDTA, 5 mM sodium butyrate, and EDTA-free protease inhibitor cocktail and incubated on ice for 10 min.

The lysate was diluted with ChIP lysis buffer lacking SDS to reduce the final SDS concentration to approximately 0.01%. Because chromatin was not sheared, cross-linked chromatin was pelleted by centrifugation and resuspended in TBS-T binding buffer containing 20 mM Tris-HCl (pH 7.4), 150 mM NaCl, and 0.05% Tween-20.

H3K27ac-associated chromatin was enriched using 5 µg anti-H3K27ac antibody (Active Motif, cat. #M0202L) and 25 µL Pierce ChIP-grade Protein A/G magnetic beads (Thermo Fisher Scientific, cat. #26162).

Magnetic beads were washed twice with TBS-T binding buffer. The washed beads were incubated with the anti-H3K27ac antibody for 1 h at 4°C with rotation. Antibody-loaded beads were then combined with the chromatin preparation and incubated overnight at 4°C with continuous rotation.

Following overnight immunoprecipitation, beads were collected magnetically and subjected to sequential washes.

Beads were washed once with low-salt wash buffer containing 150 mM NaCl, 20 mM Tris-HCl (pH 7.5), 0.1% Tween-20, and 1 mM EDTA. This was followed by two washes with high-salt buffer containing 500 mM NaCl, 20 mM Tris-HCl (pH 7.5), 0.1% Triton X-100, and 1 mM EDTA.

Beads were subsequently washed twice with stringent wash buffer containing 500 mM NaCl, 10 mM Tris-HCl (pH 8.0), 1% Triton X-100, 1% sodium deoxycholate, and 1 mM EDTA. A final wash was performed with TE buffer containing 10 mM Tris-HCl (pH 8.0) and 1 mM EDTA.

##### Elution, cross-link reversal and DNA purification

H3K27ac-enriched proximity-ligation products were eluted from the magnetic beads in buffer containing 1% SDS and 100 mM NaHCO_3_ by incubation at 65°C for 30 min. Beads were removed by magnetic separation and the chromatin-containing eluate was retained.

NaCl was added to a final concentration of 200 mM, and formaldehyde cross-links were reversed by incubation overnight at 65°C. Following reverse cross-linking, samples were treated with Proteinase K (NEB, cat. #P8107S) at a final concentration of 0.2 mg/mL in the presence of 1% SDS to digest residual protein at 65°C for 30 min. Samples were subsequently treated with RNase A (NEB, cat. #T3018L) for 15 min at 37°C. DNA was purified by phenol:chloroform:isoamyl alcohol (25:24:1) extraction. Sodium acetate (3 M, pH 5.2) was added at 0.1 sample volume, resulting in a final sodium acetate concentration of approximately 0.3 M. DNA was precipitated by addition of 2.5 volumes of 100% ethanol and incubation overnight at −20°C.Precipitated DNA was recovered by centrifugation, washed with 75% ethanol, and gently resuspended in nuclease-free water. Mechanical manipulation of the DNA was minimized throughout purification to preserve the long proximity-ligated molecules.

##### Nanopore library preparation and sequencing

Purified concatemers required no fragmentation and were converted directly into sequencing libraries with standard Oxford Nanopore ligation-based kits (SQK-LSK114 or equivalent), following the manufacturer’s end-repair, adapter ligation and clean-up steps. Libraries were quantified by Qubit and sequenced on a PromethION. Reads were typically 10–100 kb, each spanning multiple ligated fragments from an H3K27ac-enriched multi-way contact. Basecalling used Dorado(*51*) with a high-accuracy or super-high-accuracy model.

### 2. Sequencing data processing

#### Nanopore-HiChIP and Pore-C

Nanopore-HiChIP and published Pore-C data were processed with the EPI2ME Labs wf-pore-c Nextflow pipeline(*35*) (v1.3.1). Reads were digested in silico at NlaIII sites and aligned to GRCh38 (Gencode v47) with minimap2(*52*) using the map-ont preset. pore-c-py (v2.1.4) then annotated each read and resolved it into its constituent monomers; molecules containing more than 250 monomers were discarded. Valid pairwise contacts were deduplicated and aggregated into genome-wide matrices with pairtools and cooler, producing multi-resolution .mcool files at 5 kb base resolution. Separately, all multi-way (>2-fragment) contacts were extracted from each concatemer and assembled into “Chromunity” objects, which are the input for the high-order analysis below.

For allele-specific analysis, reads were assigned to parental haplotypes using WhatsHap-tagged monomer counts against the ENCODE GRCh38 phased variant call set (Table S3), summarised per read as h1_ratio = H1 / (H1 + H2). We kept only concatemers whose monomers agreed unanimously on haplotype of origin — that is, h1_ratio exactly 0 or 1 — so that no read contributes ambiguous haplotype evidence. Standard quality filters were also applied: pass-filter reads, autosomes only, blacklisted regions excluded, and at least two fragments per read.

#### Conventional HiChIP

H3K27ac HiChIP libraries were processed with the SnakeHichipTF(53) pipeline. Paired-end reads were aligned to GRCh38 (Gencode v47) and valid ligation products identified with HiC-Pro(54) (v3.1.0; bowtie2(55) v2.4.4), retaining uniquely mapped pairs with mapping quality ≥ 30, a minimum fragment length of 1 kb and interaction distances up to 2 Mb. Significant interactions were called with HiCDCplus(56) (v1.14.0) at FDR < 0.01.

#### ATAC-seq

ATAC-seq data were processed with snakePipes(57) (v3.0.0) against GRCh38 (Gencode v47). Reads were adapter- and quality-trimmed with Trim Galore (-q 20) and aligned with the DNA-mapping workflow (Bowtie2(55); mapping quality ≥ 10; PCR duplicates removed; properly paired alignments only). Fragments were restricted to 0–150 bp to select sub-nucleosomal signal, and peaks called with MACS2(58) (v2.2.7.1; q < 0.001). Coverage tracks were generated with deepTools (v3.5.6).

#### Bulk RNA-seq and GRO-seq

Total RNA-seq and GRO-seq data were processed with snakePipes(*57*) (v3.0.0) against GRCh38 (Gencode v47). Reads were aligned with STAR(*59*) (v2.7.10b) assuming a reverse-stranded library, and gene-level counts quantified with featureCounts(*60*) (-C -Q 10 --primary). Differential expression used DESeq2(*61*) (v1.42.0) with apeglm log₂ fold-change shrinkage at FDR < 0.05. Coverage tracks were generated with deepTools(*62*) (v3.5.6).

#### Allele-resolved bulk and single cell RNA-seq

Allele-specific bulk and Smart-seq2 single-cell RNA-seq were processed with an in-house Nextflow (DSL2) pipeline, wf-as-hub-burst, against GRCh38 (Gencode v47).

A specific difficulty in allele-resolved RNA-seq is reference mapping bias: reads carrying the reference allele align slightly more readily than those carrying the alternative allele, which would produce apparent allelic imbalance from a purely technical cause. We therefore aligned with STAR(59) (v2.7.11b) in WASP mode(63) (--waspOutputMode SAMtag) against each sample’s phased heterozygous variants, which re-maps every variant-overlapping read with its alleles swapped and discards reads whose mapping position changes. Reads were quality- and adapter-trimmed beforehand with Trimmomatic(64) (v0.39; ILLUMINACLIP:2:30:10:8:true, LEADING:20, TRAILING:20, SLIDINGWINDOW:5:20, MINLEN:36). We retained uniquely mapped, properly paired reads with mapping quality ≥ 20 that passed the WASP filter, and removed PCR duplicates with GATK4 (v4.6.1.0) MarkDuplicates.

Allelic counts were aggregated per gene with phASER(65) (mapping quality ≥ 255, base quality ≥ 10, genome-wide phasing), which combines evidence across all heterozygous exonic sites in a gene using phase-set-consistent assignments. This yields Allele-1 and Allele-2 count matrices across samples. Total gene expression was quantified separately with featureCounts(60) (v2.0.6).

### 3. Downstream analysis

#### Identifying high-order enhancer hubs

A set of genomic bins that appear together on the same concatemer is not by itself evidence of coordinated engagement: bins that each contact one another frequently will also co-occur by chance. Hub identification therefore tests each candidate combination against the co-occurrence expected from its own lower-order contacts.

We used Chromunity(*35*) (v0.0.2). Anchor windows were defined per chromosome from the consensus of ATAC-seq and H3K27ac ChIP-seq peaks, excluding the ENCODE blacklist and the top 1% of windows by ATAC or H3K27ac signal. Hub discovery (re_chromunity (), shave = TRUE) used a generalised linear model conditioned on four covariates that otherwise influence contact frequency: local GC content in 200-bp tiles, NlaIII restriction-fragment density, mean ATAC-seq coverage and mean H3K27ac signal (log₂ ratio). Primary parameters were cthresh = 5, k.knn = 15, k.min = 3 and peak.thresh = 0.75, with a fallback of cthresh = 3, k.knn = 5 for chromosomes on which the primary parameters failed to converge. Reads from all replicates of a condition were pooled for discovery.Candidate hubs were filtered to a cardinality, the number of distinct anchor bins, between 2 and 15. Hubs of cardinality 1 admit no multi-way comparison by construction and were discarded (9.6% of raw candidates). The upper bound 15 was imposed for computational reasons: the subset enumeration below scales as Σ ₌₁ j·C (n,j) in cardinality n at k = 7, so a small number of very large hubs would dominate the calculation entirely. In this dataset, hubs above cardinality 15 were 6.3% of candidates by count but would have generated over 99.9% of all enumerated subsets, with a single cardinality-81 outlier alone contributing more subset rows than the entire retained dataset. Such extreme outliers most plausibly reflect mapping artefacts in repetitive or unusually accessible regions rather than genuine high-order contacts.

For each retained hub, all constituent anchor bin combinations up to 7-way were enumerated (annotate (), k = 7) and tested for enrichment over the background model (synergy ()). P values were Benjamini-Hochberg corrected within each replicate separately, and a hub called significant at FDR < 0.05. Hubs significant in every replicate of a condition were designated High Confidence and used for all downstream analysis; those significant in one but not the other replicate were designated Low Confidence. Requiring significance in each replicate independently serves as a reproducibility criterion. To assess hub overlap across cell lines, two high-confidence hubs were considered overlapping if their reciprocal Jaccard index based on genomic span was ≥ 0.5.

#### Pairwise enhancer–promoter interactions

To provide a comparison class resolved at the same scale but without multi-way structure, pairwise interactions were called from the same Nanopore-HiChIP contact data with HiCDCplus(*56*) (v1.14.0) at 5 kb resolution using NlaIII/CATG restriction-fragment features, at adjusted p < 0.01, retaining only interactions reproducible across replicates. Interactions that did not overlap a High Confidence hub defined the Pairwise class. Expressed genes belonging to neither class were assigned to No_enhancer.

#### Allele-specific enhancer hub remodeling

Hub-level remodeling was assessed at two levels: overall hub contact strength and individual anchor bin participation within hubs. Hub contact strength was tested genome-wide with DESeq2(*61*). For each hub, contact counts were summed across all constituent anchor bins to obtain the hub’s total contact strength, stratified by allele. We tested whether this hub-level contact count changed between alleles using DESeq2(*61*). Hubs with FDR < 0.05 were flagged as showing significant allele-biased contact strength. This test captures whether a hub’s overall wiring is stronger on one allele, independent of which specific anchors drive that difference.

Bin-level participation was then assessed separately, addressing the distinct question of whether individual anchor bins change their relative contribution within their hub’s contact network. A hub is by construction a particular combination of bins found together on the same molecule, so asking whether that combination changed between alleles conflates several distinct events — one anchor dropping out, two anchors swapping, or the same anchors co-occurring at a different rate — into a single call. Testing each bin’s own participation instead gives an unambiguous readout: if a bin’s share of its hub’s reads changes between alleles, that bin is itself gaining or losing participation on that allele.

This bin-level test was restricted to High Confidence hubs, for which bin-resolved allelic counts could be built directly from the haplotype-tagged concatemer pool. The test is formally identical to differential exon usage, and we implemented it with DEXSeq(*38*) (v1.52.0) by treating each hub as a "gene" and its constituent anchor bins as "exons". Framing it this way controls for the hub’s overall contact strength, so that a bin is flagged only when its *relative* contribution changes, not simply because the whole locus is better covered on one allele. We used design ∼ allele + bin + condition:bin against the reduced model ∼ allele + bin, tested with testForDEU (), and obtained fold changes from estimateExonFoldChanges () with Allele-1 as reference. A bin was called lost at adjusted P < 0.05; a positive fold change denotes increased usage on Allele-2 and a negative fold change increased usage on Allele-1.

Each hub then received a single call from the pattern of its bins. Hubs with no lost bins were called Intact; hubs with at least one were called Perturbed. Among perturbed hubs, those whose lost bins all shifted toward the same allele were called Allele1-biased or Allele2-biased accordingly, and those with bins shifted in both directions were called Mixed.

#### Allele-specific transcription factor binding

Allele-specific binding (ASB) calls were taken from the ADASTRA(39) database, restricted to sites with min (FDR_ref, FDR_alt) < 0.05. Variants with mean_BAD > 1, which indicates non-diploid copy number and therefore an unreliable allelic ratio, were excluded. Binding direction was assigned from the relative effect size on allele 1 versus allele 2.

For each analysis unit, including perturbed and intact hubs and pairwise interactions, we counted three quantities: the number of phased heterozygous variants overlapping any constituent bin (n_het_SNP), how many of these were called ASB by any factor (n_ASB), and how many distinct factors showed at least one ASB variant (n_ASB_TF). The ASB rate was defined as n_ASB / n_het_SNP. Using heterozygous sites rather than region length as the denominator matters here, because regions with deeper ChIP coverage yield more testable sites and would otherwise appear to have more ASB simply for that reason.

#### Allele-specific gene expression

Allele-specific expression was tested independently in bulk and Smart-seq2 single-cell data, in both cases restricted to autosomal genes with reads per kb (RPK) ≥ 1 or count ≥ 20.

For bulk RNA-seq, we aimed to identify allele-specific expression that reflects a genuine biological signal rather than an artifact of any single dataset. Because publicly available GM12878 RNA-seq data were generated across several independently conducted ENCODE experiments, each with its own cell culture batch, library preparation, and sequencing run, differences between experiments could potentially be mistaken for allelic imbalance. We therefore combined data from such experiments while explicitly accounting for between-experiment variation, rather than averaging across them or treating all samples as interchangeable replicates.

Gene-level allelic counts from phASER(65) were retained at the level of individual biological replicates (n = 2 per experiment; 16 samples in total) and analyzed with DESeq2 using a blocked design (∼experiment + allele), with experiment of origin included as a covariate and allele as the variable of interest. This design absorbs systematic differences between experiments into the experiment term, such that the allele coefficient reflects the difference between allele 1 and allele 2 after accounting for experiment-specific variation. This is analogous to a paired comparison in which each experiment provides an internal allele 1- versus-allele 2 contrast. Preserving the original biological replicates was important for this design: variation among biological replicates within the same experiment and allele allows DESeq2 to estimate the residual variability against which the allelic effect is evaluated. Genes were called allele-biased at FDR < 0.05 and |log₂ fold-change| ≥ 1.

For single-cell data, allelic counts were summed across cells and tested with Wilcoxon test compared per-cell Allele-1 and Allele-2 counts across all cells with non-zero allelic coverage and was Benjamini-Hochberg corrected.

#### Transcriptional bursting kinetics

Bursting parameters were inferred separately for each allele of each gene from the Smart-seq2 allelic counts using txburst(*42*), which fits the Poisson-Beta formulation of the two-state promoter model by maximum likelihood and returns kon, koff and ksyn with profile-likelihood confidence intervals. Burst frequency was taken as kon and burst size as ksyn/koff. Allelic differences were expressed as log₂ fold changes between haplotypes. Perturbed-hub genes with |log₂FC| > 0.5 in either parameter were designated dynamic hubs and those below this threshold static hubs.Because these parameters are model estimates rather than direct measurements, we checked that they track features visible in the raw counts. For each gene we computed two statistics with no model fitted: the difference between alleles in the fraction of cells with zero counts (silent cells), which reflects how often the gene fires, and the difference in mean count among active expressing cells, which reflects how much it produces per firing.

We also tested whether the relationship between the two parameters could be an artefact of estimation precision, since more highly expressed genes are measured more accurately. Pairwise-regulated and static-hub genes were matched to dynamic-hub genes on log expression level and log allelic read depth by propensity-score nearest-neighbour matching without replacement, with a caliper of 0.2 standard deviations of the logit propensity score. The Pearson correlation between allelic burst frequency and burst size was then recomputed on 1,000 bootstrap resamples of each matched control set, and the observed dynamic-hub correlation compared against those distributions.

#### Buffering Index

To express how much kinetic variation fails to reach expression, we decomposed the variance of allelic mean expression into contributions from burst frequency (f), burst size (b) and their covariance (Cov). Since mean expression is the product of the two parameters, reciprocal changes cancel in the covariance term, and the Buffering Index is the fraction of total kinetic variance cancelled that way:

BI = −2 Cov (Δlog f, Δlog b) / (Var (Δlog f) + Var (Δlog b))

A value of 0 means the two parameters vary independently, so kinetic changes pass through to expression; a value of 1 means they cancel exactly, so expression is unchanged. Confidence intervals were obtained by gene-level bootstrap resampling with 1,000 replicates.

#### Dosage sensitivity and disease variant analysis

All analyses in this section were run separately for each cell line, since hubs are called from each cell line’s own data and are not pooled.

Haploinsufficiency was assessed with four independently derived measures: haploinsufficiency score from the dosage sensitivity map (haploinsufficiency index)(*1*), the gnomAD loss-of-function intolerance probability(*6*), the ExAC loss-of-function z score(*43*), and sHet, an estimate of the selection coefficient against heterozygous loss of function(*44*) (Table S3). These differ in derivation and scale but all increase with intolerance to losing one copy. Experimentally curated haploinsufficient genes were taken from ClinGen(*45*). Enrichment of ClinGen tier 3 haploinsufficient genes was tested with two-by-two Fisher’s exact tests, hub versus pairwise and hub versus no-enhancer. Comparisons used all High Confidence hub genes, since the question is whether a gene is hub-regulated at all rather than the direction of its allelic imbalance.GWAS-associated variants(*46*) and ClinVar(*47*) pathogenic and likely pathogenic variants were intersected with hub bin and pairwise anchor coordinates. Densities were expressed per kilobase of regulatory sequence, since hub regions and pairwise anchors differ substantially in size.

cis-eQTL summary statistics came from the EBI eQTL Catalogue(*12*, *48*) (GTEx v8, study QTS000015), using tissues matched to each cell line: whole blood and EBV-transformed lymphocytes for GM12878, and lung for IMR90. eQTL variants were assigned to hub or pairwise regions by positional overlap and absolute effect sizes compared between classes. Because eQTL effect size depends strongly on minor allele frequency and on distance to the transcription start site, the comparison was repeated with adjustment for both.

### Functional enrichment

Enrichment among hub-target genes was tested with Enrichr(*66*) (v3.4; Fisher’s exact test) against GO Biological Process 2025(*67*) and DisGeNET(*68*), reported at adjusted P < 0.05.

### Statistics and reproducibility

Unless otherwise stated, two-group comparisons used two-sided Wilcoxon rank-sum tests, categorical comparisons used Fisher’s exact tests, and multiple testing was controlled by the Benjamini-Hochberg procedure at FDR < 0.05. All analyses were performed independently in GM12878 and IMR90.

## Acknowledgements

We thank Dr. Jeffrey Milbrandt and Dr. Sheng Chih Jin for insightful discussions and continuous support of this project. We thank all members of our department and institutes for fostering a collaborative and supportive research environment.

## Authors’ contributions

J.T., Y.S., and T.W. conceived and designed the study. J.T. led the development of the computational framework, J.T., Y.W., and C.P. performed the computational, allele-specific, genomic, and transcriptomic analyses. M.F.S., C.F., C.M., X.C., and R.S.F. contributed to the method development, generation, and processing of Nanopore-HiChIP data. R.H. contributed to genomic and allele-specific analyses. J.T., T.W. and Y.S. interpreted the results and developed the overall mechanistic model. T.W. and Y.S. supervised the project. J.T., T.W., and Y.S. wrote the manuscript, with contributions from all authors. All authors reviewed and approved the final manuscript.

## Data availability

Nanopore-HiChIP data generated from GM12878 and IMR90 cells are available in the Gene Expression Omnibus (GEO) under accession number GSE346397. Pore-C data for GM12878 were obtained from GSE149117. ATAC-seq data for GM12878 and IMR-90 were obtained from GSE170245 and GSE169767, respectively. H3K27ac, H3K4me1, and H3K4me3 ChIP-seq data for GM12878 and IMR-90 were obtained from ENCODE portal: ENCSR000AKC and ENCSR002YRE, ENCSR000AKF and ENCSR831JSP, and ENCSR057BWO and ENCSR087PFU, respectively. GRO-seq data for GM12878 and IMR90 were obtained from GSE60454 and GSE43070, respectively, and PRO-seq data were obtained from GSE214304 and GSE105936, respectively. Bulk poly(A) RNA-seq data were obtained from GSE78550, GSE78551, GSE78552, GSE78553, GSE78554, GSE78555, GSE90276 for GM12878 and from GSE90257, GSE90262, GSE90263 for IMR90. Single-cell Smart-seq2 RNA-seq data for GM12878 and IMR90 were obtained from GSE240114 and GSE115301, respectively. All datasets and their corresponding accession numbers are provided in Table S4.

## Code availability

The allele-specific analysis pipeline is available at https://github.com/YidanSunResearchLab/wf-as-hub-burst.git.

---

**Figure S1.**
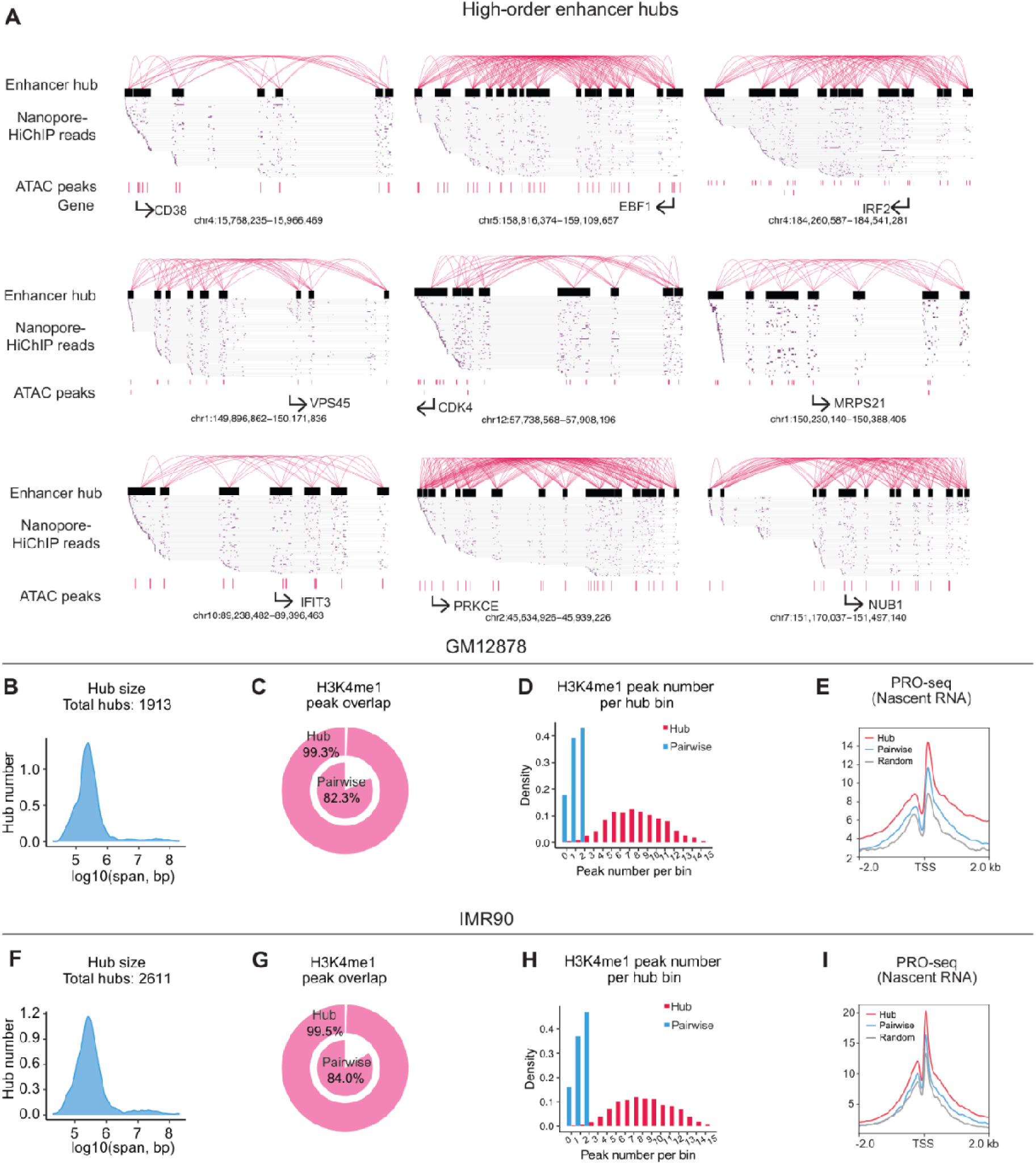
Characterization of high-order enhancer hubs. **(A)** Nine additional representative enhancer hubs in GM12878, including the CD38, EBF1 and IRF2 loci previously identified as multi-way regulatory hubs by independent methods. Arcs show hub contacts; each grey line is a single Nanopore-HiChIP read; ATAC-seq peaks are shown beneath. **(B, F)** Genomic span of enhancer hubs in GM12878 (B) and IMR90 (F). **(C, G)** Fraction of hub bins (outer ring) and pairwise enhancer interactions (inner ring) overlapping an H3K4me1 peak, in GM12878 (C) and IMR90 (G). **(D, H)** Number of H3K4me1 peaks per bin for hub bins (red) and pairwise interactions (blue), in GM12878 (D) and IMR90 (H). **(E, I)** Metagene profiles of PRO-seq nascent transcription around transcription start sites for hub-target, pairwise-regulated and random gene sets, in GM12878 (E) and IMR90 (I).

**Figure S2.**
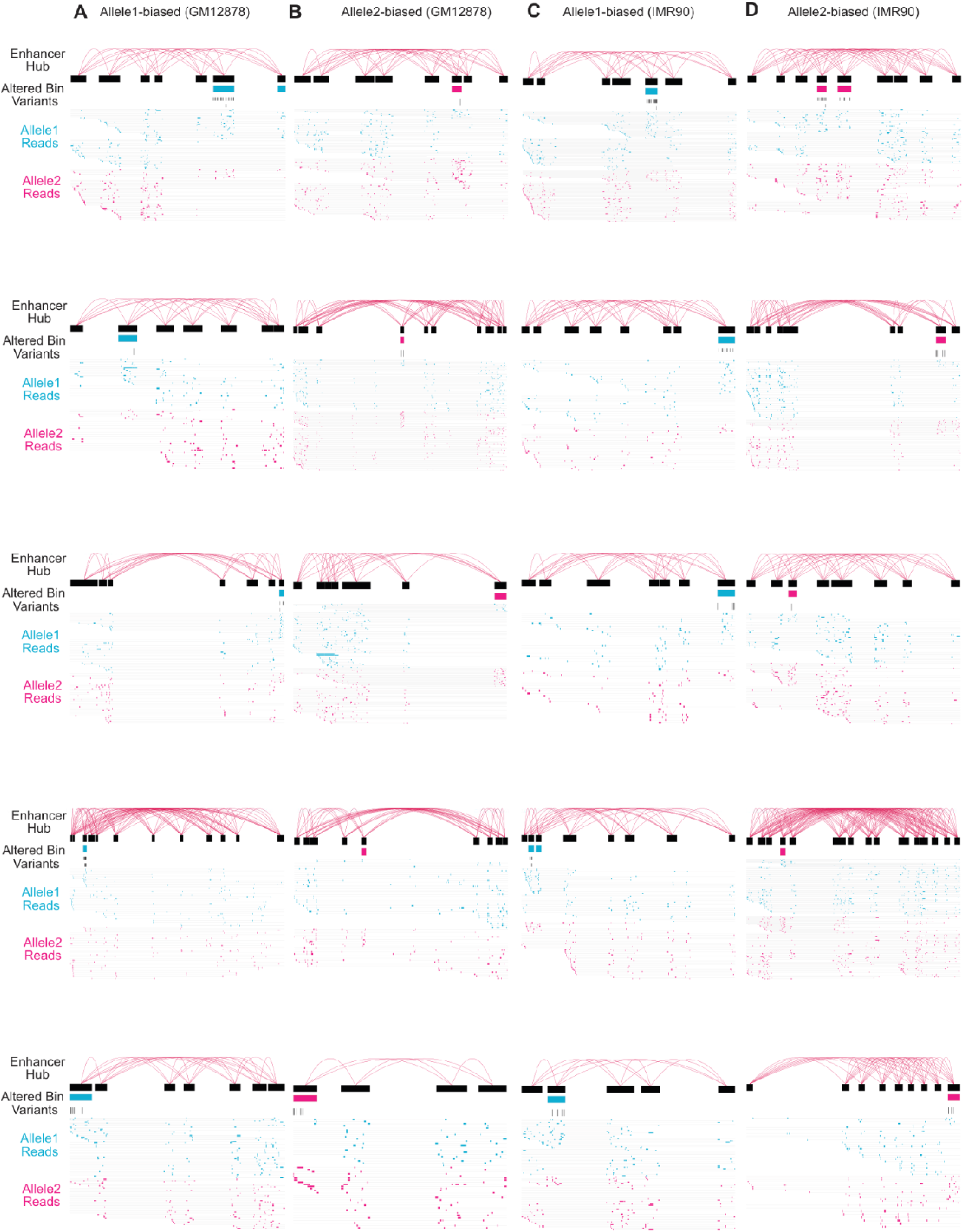
Representative examples of allele-specific enhancer hub remodeling. Twenty representative allele-biased hubs: allele 1–biased **(A)** and allele 2–biased **(B)** in GM12878, and allele 1–biased **(C)** and allele 2–biased **(D)** in IMR90. For each, arcs show hub contacts, the altered bin is highlighted, heterozygous variants within it are marked, and individual reads assigned to allele 1 (blue) and allele 2 (red) are shown below.

**Figure S3.**
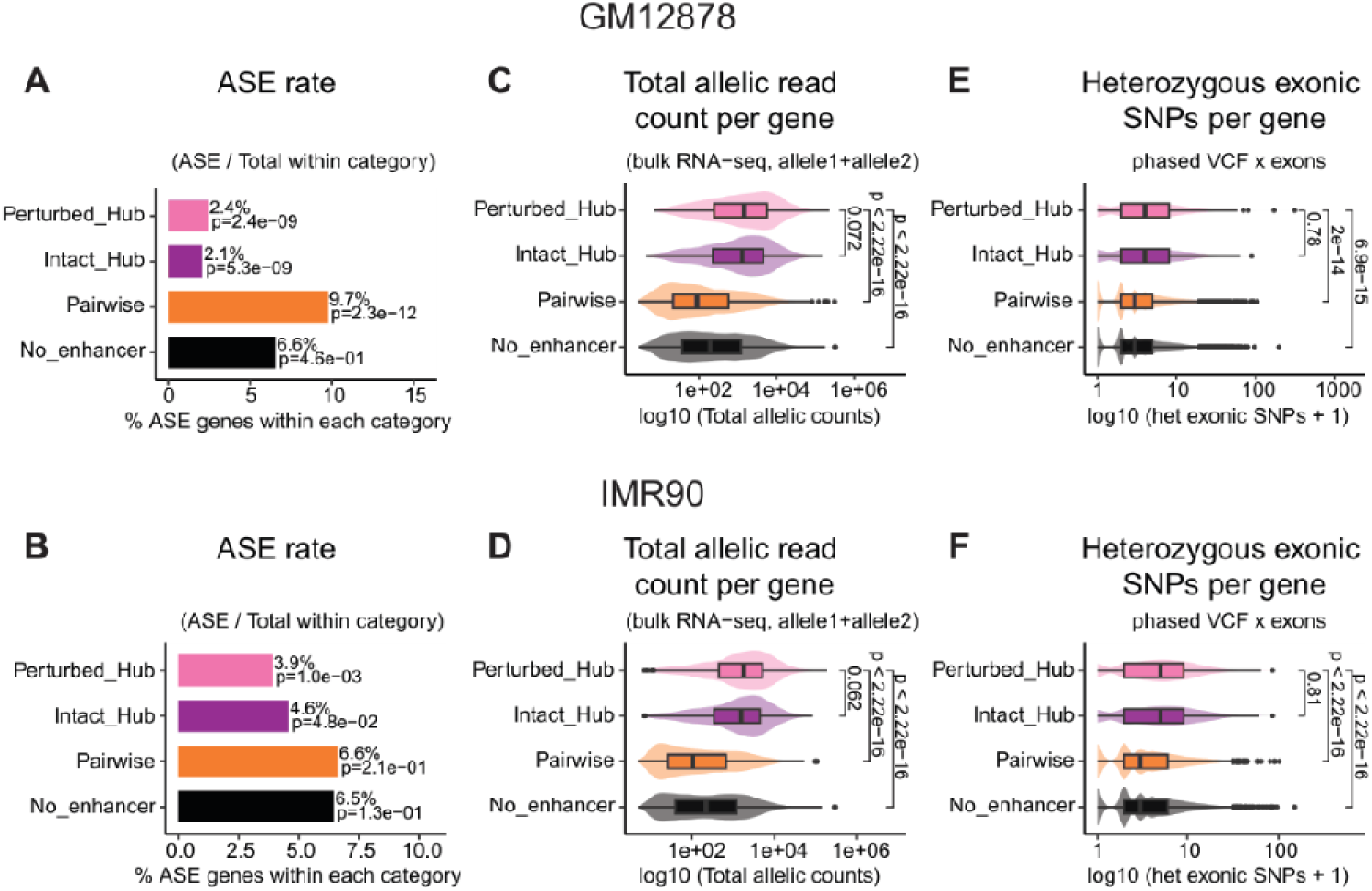
Depletion of allele-specific expression at hub-target genes is not a power limitation. **(A, B)** ASE rate within each regulatory class — the fraction of genes in that class reaching significance — in GM12878 (A) and IMR90 (B). This complements the composition view in Fig. 3C, D by controlling for the different numbers of genes in each class. Fisher’s exact test versus all other classes. **(C, D)** Total allelic read counts per gene by regulatory class in GM12878 (C) and IMR90 (D). **(E, F)** Number of heterozygous exonic variants per gene by regulatory class in GM12878 (E) and IMR90 (F). Read depth and heterozygous site count are the two principal determinants of power to detect allelic imbalance. Hub-target genes are equal or higher on both, so on technical grounds they should yield more ASE calls than the comparison classes, not fewer. ns, not significant; **P < 0.01; ****P < 0.0001, two-sided Wilcoxon rank-sum test is used.

**Figure S4.**
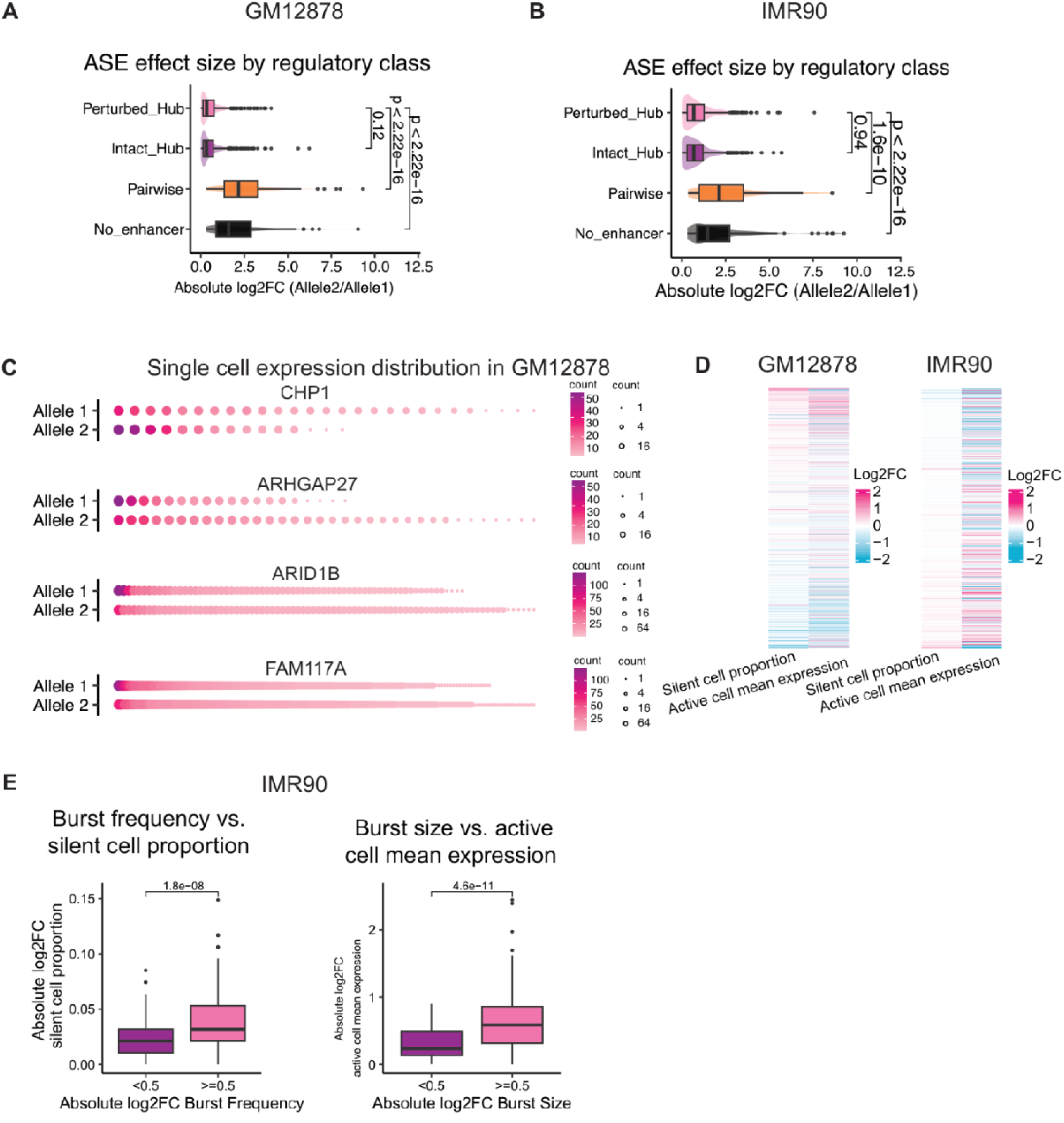
Allele-resolved single-cell kinetic analyses. **(A, B)** Absolute allelic fold change in mean expression from single-cell data, by regulatory class, in GM12878 (A) and IMR90 (B). The single-cell measurements reproduce the bulk RNA-seq result, confirming that balanced output at hub-target genes is not an averaging artefact of bulk measurement. Two-sided Wilcoxon rank-sum test is used. **(C)** Allele-resolved single-cell expression distributions for three representative genes in GM12878, plotted as in Fig. 4A. **(D)**Allelic differences in the two raw distributional features, the fraction of silent cells and mean expression among active cells, for individual perturbed-hub genes in GM12878 (left) and IMR90 (right), ordered by silent-cell fraction difference. **(E)** Fitted kinetic parameters track the raw single-cell distributions in IMR90, as in Fig. 4F.

**Figure S5.**
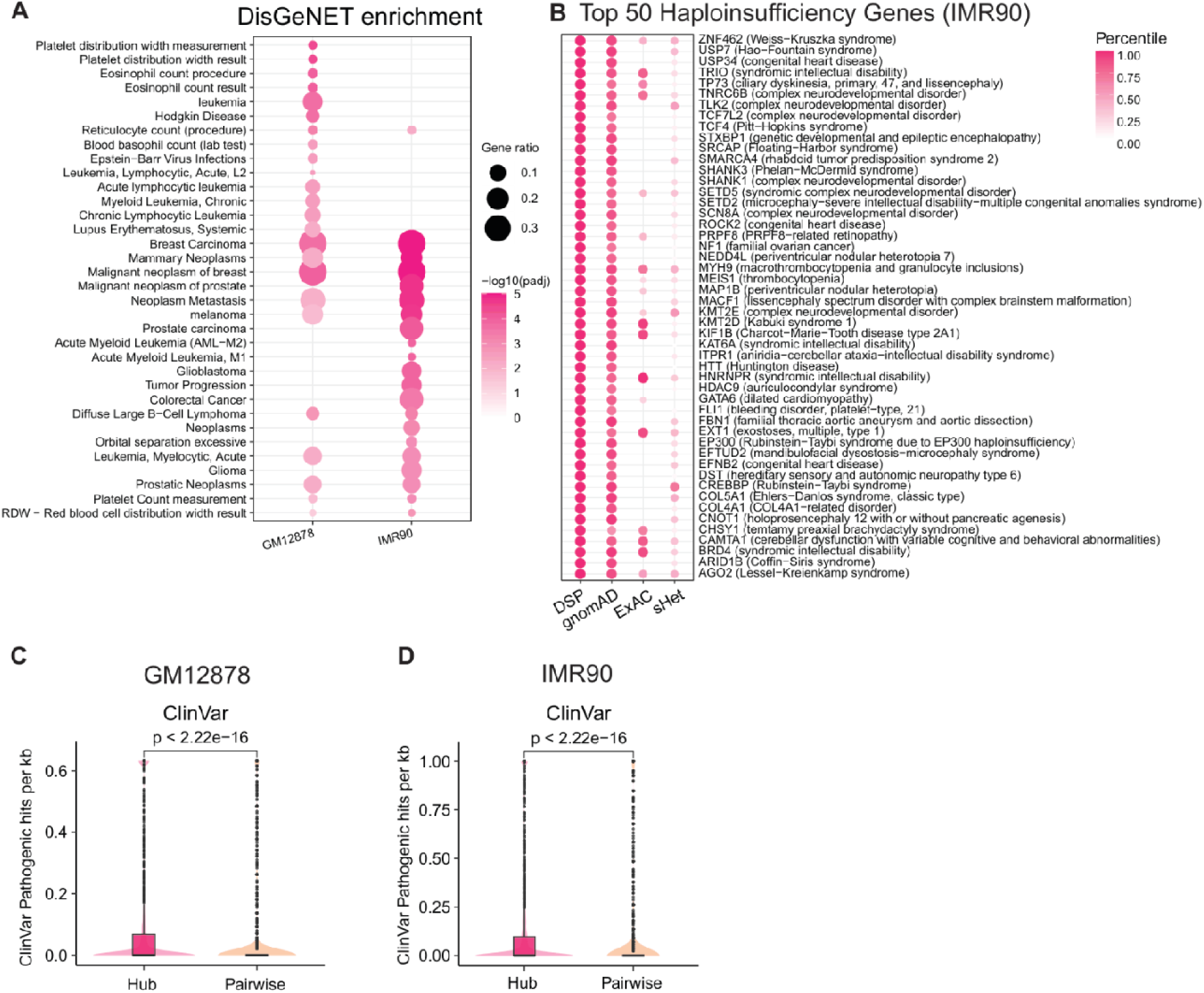
Additional disease and dosage-sensitivity analyses. **(A)** DisGeNET disease-term enrichment for hub-target genes in each cell line. Enriched terms follow cell identity, with leukaemias and lymphomas in GM12878 and solid tumours in IMR90. **(B)** The 50 most haploinsufficient hub-target genes in IMR90 with their associated diseases, plotted as in Fig. 5G. **(C, D)** Density of ClinVar pathogenic and likely pathogenic variants per kilobase of regulatory sequence in hub regions versus pairwise enhancer anchors, in GM12878 (C) and IMR90 (D). Two-sided Wilcoxon rank-sum test is used.

